# A mechanism for ligand gated strand displacement in ZTP riboswitch transcription regulation

**DOI:** 10.1101/521930

**Authors:** Eric J. Strobel, Luyi Cheng, Katherine E. Berman, Paul D. Carlson, Julius B. Lucks

## Abstract

Cotranscriptional folding is an obligate step of RNA biogenesis that can guide RNA structure and function by forming transient intermediate folds. This is especially true for transcriptional riboswitches in the which the formation of ligand-dependent structures during transcription regulates downstream gene expression. However, the intermediate structures that comprise cotranscriptional RNA folding pathways and the mechanisms that enable transit between them remain largely unknown. Here we determine the series of cotranscriptional folds and rearrangements that mediate antitermination by the *Clostridium beijerinckii pfl* riboswitch in response to the purine biosynthetic intermediate ZMP. We uncover sequence and structural determinants that modulate a regulatory RNA strand displacement reaction and identify biases within natural ZTP riboswitch sequences that promote on-pathway folding. Our findings establish a mechanism for ZTP riboswitch antitermination and suggest general strategies by which nascent RNA molecules can navigate cotranscriptional folding pathways efficiently.

## Introduction

The coupling of transcription and RNA folding is ubiquitous to RNA biogenesis^1^. Nascent RNA folding is directed by the 5’ to 3’ polarity of transcription and the typically slower rate of the nucleotide addition cycle relative to base pair formation^2–4^. Consequently, cotranscriptional RNA folding favors the formation of local structures that can pose energetic barriers to the formation of long-range interactions^5–7^. The tendency of RNA molecules to enter such kinetic traps as they fold during transcription is thought to be the basis for gene regulation by riboswitches, non-coding RNAs that adopt alternate conformations to control gene expression in response to chemical ligands^8–11^. The identification of riboswitch ligands has consistently revealed diverse roles for RNA molecules in cellular physiology and illustrated the capacity of RNA for precise chemical recognition^12,13^. In addition, riboswitches have found diverse utility as antibiotic targets^14^, diagnostic biosensors^15,16^, and as imaging tools^17^.

The general architecture of a riboswitch comprises a ligand-sensing aptamer domain and an ‘expression platform’ that directs a regulatory outcome based on ligand occupancy^18^. In some cases, these domains share an overlap sequence that participates in mutually exclusive aptamer and expression platform structures that either block or allow downstream gene expression^9–11^. Cotranscriptional RNA folding is integral to the function of riboswitches that regulate transcription because ligand recognition must occur within a limited window ^19,20^. Ligand binding is thought to kinetically trap the aptamer fold, effectively sequestering the overlap sequence and preventing the expression platform from forming^21–24^. While crystal structures of diverse ligand-bound riboswitch aptamers have provided a detailed understanding of ligand-aptamer complexes^12^, the sequence of folding intermediates that mediate riboswitch function has only been described in a handful of cases^21,22,25^. Thus, understanding how ligand-dependent aptamer stabilization modulates expression platform folding to control riboswitch function during transcription remains a major goal.

Here we investigated how a riboswitch that senses the purine biosynthetic intermediate 5-aminoimidazole-4-carboxamide riboside 5’-triphosphate (ZTP)^26^ mediates transcription antitermination. The identification of the ZTP riboswitch uncovered a mechanism by which ZTP and its monophosphate derivative ZMP (Fig. 1a) function as bacterial alarmones for deficiency in the cofactor 10-formyl-tetrahydrofolate (10f-THF) to signal the expression of genes associated with 10f-THF biosynthesis^26,27^. The ZTP aptamer comprises a helix-junction-helix motif (P1-J1/2-P2) and a small hairpin (P3) that are separated by a variable length linker but interact to form the ZTP binding pocket^26,28–31^ (Fig. 1b). In the *Clostridium beijerinckii* (*Cbe*) *pfl* ZTP riboswitch, the intrinsic terminator overlaps the aptamer such that the 5’ terminator stem comprises the P3 hairpin and forms a pseudoknot with J1/2^26^ (Fig. 1b). Crystallographic studies of several ligand-bound ZTP aptamers revealed an extensive network of contacts between the aptamer subdomains that could stabilize the aptamer against transcription termination^28–30^. However, the ZMP-dependence of these interactions during transcription remains unclear and the complexity of the aptamer sub-domain interface obscures the precise mechanism through which aptamer state drives antitermination.

**Figure 1.**
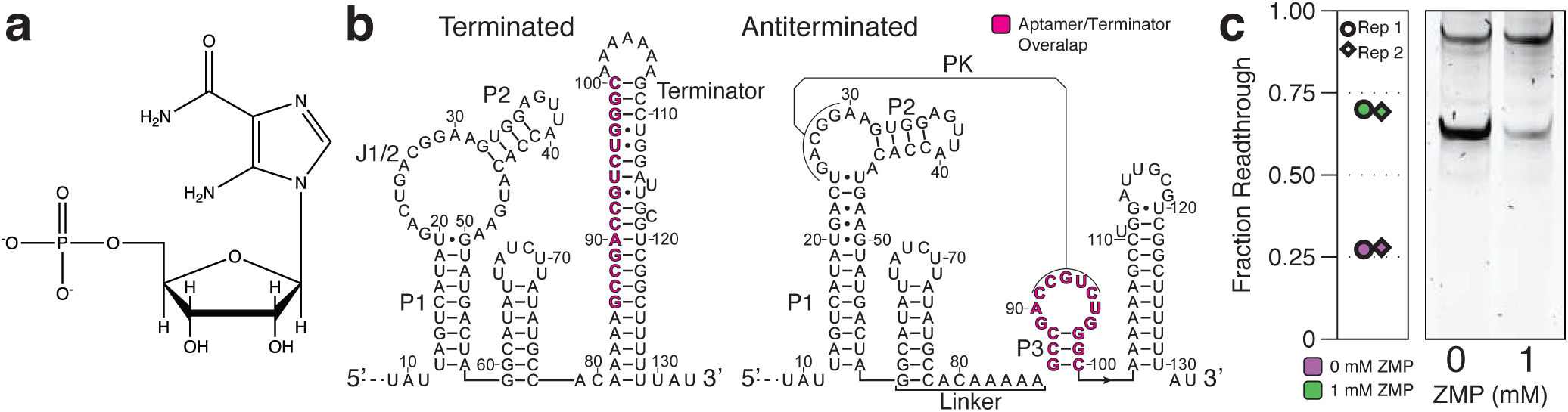
Overview of the *Clostridium beijerinckii pfl* ZTP riboswitch. **(a)** Chemical Structure of ZMP. **(b)** Secondary structure models of the terminated and antiterminated folds derived from covariation analysis^26^ and x-ray crystallography^28–30^. Three paired regions (P1-P3), the terminator hairpin, the junction between P1 and P2 (J1/2), and the linker between the P1-J1/2-P2 and P3 aptamer subdomains are annotated. The aptamer/terminator overlap sequence is highlighted in magenta. **(c)** *In vitro* transcription in the presence and absence of 1 mM ZMP. Reaction conditions match those used for cotranscriptional SHAPE-Seq. Fraction read through values are quantified from the gel image. n=2 independent biological replicates are shown.

**Figure 1-source data.** Source data for Figure 1 are available in the Northwestern University Arch Institutional Repository (https://doi.org/10.21985/N20F4H).

In this work, we first characterize ligand-dependent folding pathways for the Cbe *pfl* ZTP riboswitch using high-throughput nascent RNA structure probing. Our analysis revealed three key features of *pfl* aptamer folding pathway: 1) the aptamer fold is preceded by a transient intermediate, 2) organization of the overall aptamer tertiary fold is ZMP-independent but ZMP binding stabilizes a network of discrete tertiary structure interactions, and 3) ZMP binding must occur within a short window of 7-15 nucleotide addition cycles. We then apply a combinatorial mutagenesis strategy to uncover how ZMP binding mediates transcription antitermination by identifying the pathway and sequence determinants of terminator hairpin nucleation and folding. Combined with our structural measurements, our functional analysis pinpoints the ligand-dependent inaccessibility of P3 to terminator nucleation as the principal determinant of antitermination. Furthermore, an analysis of diverse ZTP riboswitch sequences revealed that P3 has context-dependent sequence preferences that may avoid off-pathway states to promote efficient aptamer folding in the short window for ligand binding. Overall, our findings reveal the mechanism of *pfl* ZTP riboswitch antitermination and suggest general folding strategies that could guide cotranscriptional structure formation in diverse RNA molecules.

## Results

### A transient intermediate fold precedes nascent ZTP aptamer folding

To uncover ZTP riboswitch folding intermediates in the presence and absence of ZMP (Fig. 1c), we used cotranscriptional Selective 2’-Hydroxyl Acylation analyzed by Primer Extension and Sequencing (SHAPE-Seq), which couples high-throughput RNA structure probing (reviewed in^32^ and^33^) with roadblocked *in vitro* transcription to chemically probe an array of nascent RNA transcripts^22,34^ (Fig. 2). Initial *pfl* aptamer folding is defined by the ZMP-independent formation of an intermediate hairpin (IH1) from nts 12-25 at transcript length 45 (Fig. 2a, b). IH1 persists through transcripts ~45 to ~70, during which P2 folds as indicated by high reactivity at nts 38-40 (Fig. 2a). At transcript length ~70 a gradual decrease in IH1 loop reactivity inversely correlates with increased reactivity throughout J1/2 when P1 nucleotides 50-58 emerge from RNA polymerase (RNAP) and drive refolding into the mutually exclusive P1 fold that comprises the first aptamer subdomain (Fig. 2c, e). SHAPE probing of equilibrated *pfl* intermediates revealed a sharp delineation between the IH1 and P1 structures across transcripts 55 and 56, suggesting that IH1 is thermodynamically favorable until P1 can form (Supplementary Fig. 1). Because cotranscriptional SHAPE-Seq probes RNAs within roadblocked elongation complexes, IH1 folding may be enabled by transcription arrest. However, because the timescale of formation of local RNA structures is typically orders of magnitude faster than nucleotide addition by bacterial RNAPs^1,35^, the persistence of IH1 for at least 35 nt addition cycles suggests that IH1 folds even during uninterrupted transcription. Minimum free energy structure prediction by RNAstructure’s Fold method^36^ indicated that ~50% of ZTP riboswitch sequences from 532 fully sequenced bacterial genomes^26^ can form an IH1-like structure at least as favorable as *Cbe* IH1 (Supplementary Fig. 2a). Structure prediction using randomized sequences that were constrained by natural nucleotide frequency approximated the distribution of free energies for natural ZTP riboswitch sequences (Supplementary Fig. 2b). These analyses combined with probing data suggest that transient formation of a transient intermediate hairpin prior to P1 folding is a feature of some ZTP riboswitches that is favored by the sequence constraints of the aptamer and may explain the tendency of some ZTP aptamers to misfold under equilibrium conditions^26,29^.

**Figure 2.**
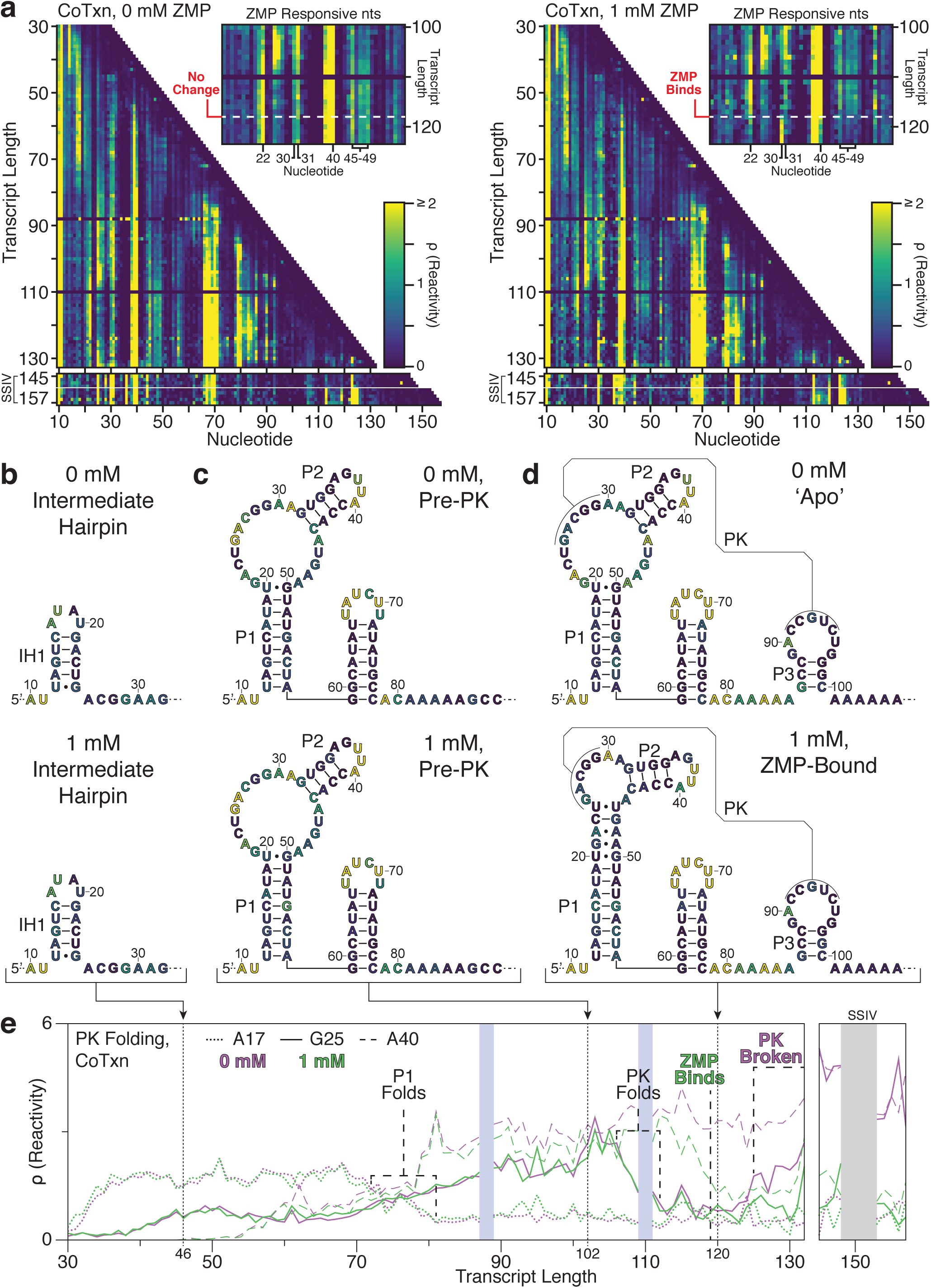
*C. beijerinckii pfl* ZTP riboswitch folding intermediates. **(a)** Cotranscriptional SHAPE-Seq reactivity matrix for the *Clostridium beijerinckii pfl* ZTP riboswitch with 0 mM and 1 mM ZMP. ZMP-responsive nts are highlighted in reactivity matrix cut-outs. The highly reactive unstructured leader (nts 1-9) is not shown and the absence of data for transcripts 88 and 110 is due to ambiguous alignment of 3’ end sequencing reads. Reactivity spectra for transcripts 145-148 and 153-157 are from a separate experiment using terminal biotin-streptavidin roadblocks and Superscript IV reverse transcriptase (RT) to resolve elevated RT stalling in transcripts beyond the terminator. **(b)** Intermediate hairpin (IH1) secondary structure identified by manual analysis of reactivity values and refined with minimum free energy structure predictions of the leader region. Nucleotides are colored by the reactivity of transcript length 46 from (a). **(c)** Secondary structure of P1-J1/2-P2 and the linker region based on the *pfl* RNA motif structure^26^. Nucleotides are colored by the reactivity of transcript length 102 from (a). **(d)** Apo and ZMP-bound secondary structure as determined by manual SHAPE-refinement of the consensus structure (Apo)^26^ or by the ZTP aptamer crystal structures^28–30^, respectively. Nucleotides are colored by the reactivity of transcript length 120 from (a). Nucleotides at the RNA 3’ end of RNA structures that lie within the RNAP footprint are omitted in panels (b-d). **(e)** ZTP riboswitch folding as depicted by reactivity trajectories for nucleotides A17 (within IH1 loop or P1 stem, dotted lines), G25 (within J1/2 or the pseudoknot, solid lines), and A40 (within P2 loop, dashed lines) from (a). Trajectories are shown for both the absence (purple) and presence (green) of 1mM ZMP. Notable structural changes are identified by vertical dashed lines and the position of critical intermediate structures in (b-d) are shown by vertical dotted lines. Shaded boxes indicate the regions of poor alignment described in (a). Results shown are from one representative of n=3 independent biological replicates (a, upper matrices; e) or n=2 independent biological replicates (a, lower matrices, e); Replicate data and correlations are shown in Supplementary Figure 5.

**Figure 2-source data.** Source data for Figure 2 are available in the Northwestern University Arch Institutional Repository (https://doi.org/10.21985/N20F4H).

### ZTP aptamer pseudoknot folding is independent of ZMP Binding

Following rearrangement of IH1 to form P1, we next observed the folding of a hairpin from nts 59-78 and an adjacent 7 nt unstructured region that comprise a linker between P1 and P3 (Fig. 2c). ZMP-independent pseudoknot folding is then observed as a decrease in reactivities at nts 25-29 across transcript lengths 106-112, when nts 92-95 are expected to have emerged from RNAP given the ~14 nt footprint of *E. coli* RNAP on nascent RNA^37^ (Fig. 2e). Equilibrium refolding also revealed abrupt pseudoknot formation at transcript 95, when complete pairing between nts 25-29 and 91-95 first becomes possible (Supplementary Fig. 1). Together, these data indicate that the pseudoknot can fold before the P3 stem forms, though ZMP-dependent reactivity changes are only observed after the entire P3 stem has emerged from RNAP around transcript length 117 to form the complete aptamer structure (Fig. 2a, d, e). Consistent with this observation, SHAPE probing of equilibrium refolded RNAs reveals ZMP-dependent reactivity changes precisely when the P3 stem is expected to fold at transcripts 99 and 100 (Supplementary Fig. 1). Furthermore, mutations that disrupt and restore the pseudoknot disrupt and restore observed pseudoknot folding and ZMP-binding, respectively (Supplementary Figs. 3 and 4). The requisite folding of P3 prior to the observation of any ZMP-dependent reactivity differences is consistent with the observation that the P3 stem stacks with the highly conserved G97 to form the floor of the ZMP binding pocket^28–30^.

### ZMP binding is associated with coordinated multi-point aptamer stabilization

Comparison of nascent ‘apo’ and ‘holo’ intermediate transcripts revealed ZMP-responsive nucleotides that are consistent with equilibrium measurements made by in-line probing^26^ and can be categorized as P1-associated or pseudoknot-contacting based on comparison to the ZMP-bound crystal structures^28–30^ (Fig. 3 and Supplementary Fig. 6). In both cotranscriptional and equilibrium SHAPE-Seq experiments we observe a coordinated ZMP-dependent reactivity decrease across nts G21, A22, A45 and nts 47-49 at transcript lengths 117 and 100, respectively(Fig. 3a and Supplementary Fig. 6a). These signatures suggest that formation of a primarily non-Watson-Crick (WC) helical extension of P1 that was observed in the ZMP-bound crystal structures is ZMP-dependent^28–30^. The observed ZMP-dependent reactivity changes in P1 are coordinated with reactivity changes at nts A30, A31, U39, and A40, which localize to either edge of the pseudoknot^26,28–30^ (Fig. 3b, Supplementary Fig. 6b). Decreased reactivity at U39 and the highly conserved A40 is consistent with formation of a Type I A-minor interaction between A40 and the J1/2:P3 pseudoknot^26,28–30^, whereas a ZMP-dependent increase at A31 may be due to formation of a bulge upon stacking of A30 and the G26:C94 pseudoknot base pair^30^. Disruption of the pseudoknot renders each of the above nucleotides ZMP-non-responsive (Supplementary Fig. 6). The ZMP-dependent stabilization of contacts between A40/A31 and each edge of the pseudoknot suggests that ZMP-binding coordinates a P2 conformation that favors winding of the non-WC P1 base pairs, which in turn may favor the formation of the P1:P3 ribose zipper^26,28–30^. While we observe these features as a coordinated reactivity change across many nucleotides, our data cannot determine whether these changes happen in concert or in a series of folding events.

**Figure 3.**
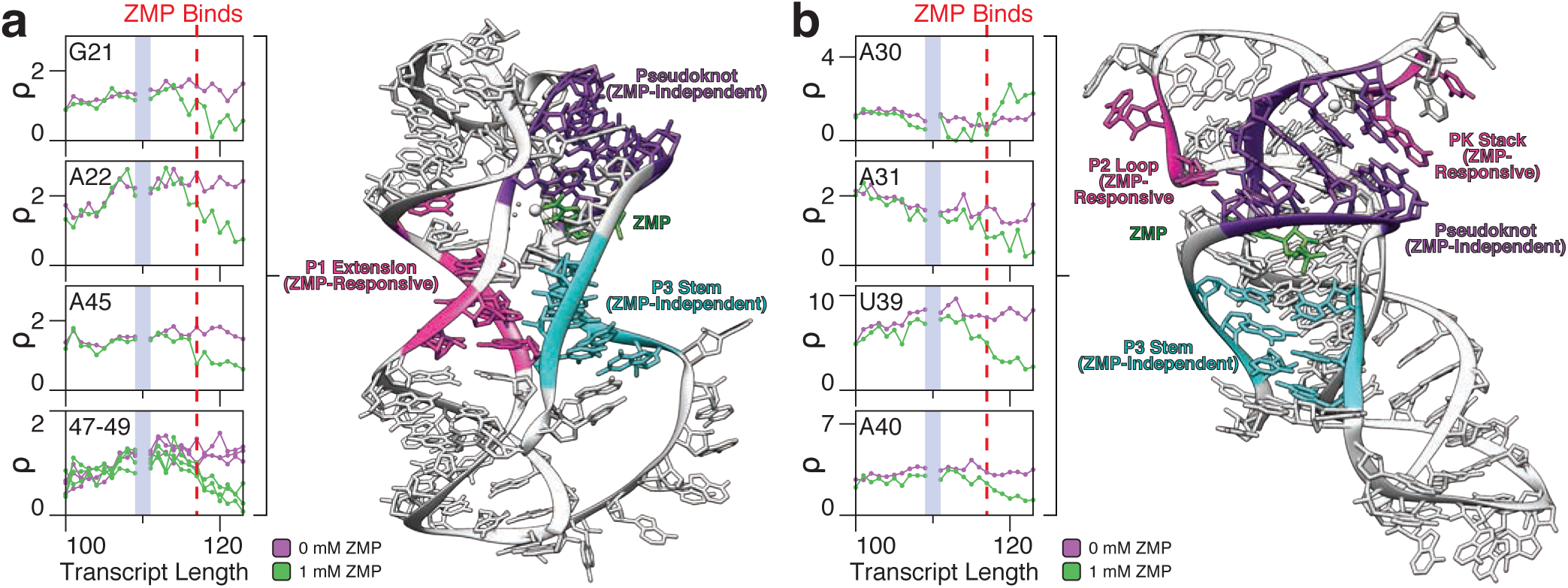
ZMP-responsive nucleotides reveal multipartite aptamer stabilization. ZMP-responsive nucleotides within the *Cbe* ZTP riboswitch, including **(a)** non-canonical P1 base pairs and **(b)** the nucleotides that interact with the pseudoknot, are shown in the absence (purple) and presence (green) of 1mM ZMP. Data are from Figure 2a (WT *Cbe* cotranscriptional SHAPE-Seq replicate 1). The vertical red dashed lines indicate the transcript length at which ZMP binding is inferred to occur based on bifurcation of the reactivity trajectories. The crystal structure of the *Thermosinus carboxydivorans* ZMP riboswitch (PDB: 4ZNP)^30^ is highlighted to show ZMP-stabilized P1 nuceotides (a) and ZMP-dependent pseudoknot-interacting nts (b). ZMP-responsive nts are highlighted in magenta, the pseudoknot is highlighted in purple, the P3 stem is highlighted in cyan, and ZMP is highlighted in green. Results shown are from one representative of n=3 independent biological replicates.

**Figure 3-source data.** Source data for Figure 3 are available in the Northwestern University Arch Institutional Repository (https://doi.org/10.21985/N20F4H).

### The ZMP-bound aptamer fold persists beyond antitermination when cotranscriptionally folded

We observe the primary termination site at nt 132, only ~15 nts after ZMP binding can occur (Fig. 2e). The first observed signature of terminator folding is a gradual increase in reactivity at the J1/2 pseudoknot nts 25-29 from transcript lengths ~125-132 in the absence of ZMP, suggesting that pseudoknot disruption can begin shortly after nucleotides within the 3’ terminator stem have emerged from RNAP (Fig. 2a, e). In contrast, sustained low reactivity at J1/2 in the presence of ZMP suggests that the pseudoknot remains stable (Fig. 2a,e). The sole deviation from this trend is the primary termination site at transcript length 132 where transcripts that terminate naturally or as a consequence of roadblocking at the termination site lead to increased J1/2 reactivity even in the presence of ZMP (Fig. 2e). The final riboswitch fold was obscured due to Superscript III reverse transcriptase stalling in transcripts beyond the terminator. To overcome this, we performed a separate experiment that enriched for transcript lengths 145 and 155 and used Superscript IV to improve full-length cDNA yield (Fig. 2a). Consistent with the observation that different RTs have distinct adduct detection biases, SSIV reactivity measurements were sparse but in qualitative agreement with measurements made using SSIII^38,39^ (Fig. 2a). The reduced reactivity at G25, A31, and A40 observed with SSIII in the presence of ZMP after transcript length 117 is recapitulated across all post-termination site lengths with SSIV, suggesting that the ZMP-bound ON state persists, whereas high reactivity at G25, A31, A40 in the absence of ZMP is consistent with pseudoknot disruption (Fig. 2a, e). In contrast, under equilibrium refolding conditions the pseudoknot cannot outcompete the terminator fold regardless of the presence of ZMP (Supplementary Fig. 1). This is similar to our finding for the fluoride riboswitch^22^ and suggests that the ZTP riboswitch antiterminated fold is only accessible by a cotranscriptional folding regime.

### Optimal *pfl* aptamer sequence variants balance pseudoknot and terminator pairing states

While cotranscriptional SHAPE-seq identifies intermediate nascent RNA structures, it does not reveal how RNA molecules transit between these structures. To ask how specific nucleotide interactions mediate ZTP riboswitch folding and antitermination, we developed a combinatorial mutagenesis strategy to perturb RNA folding (Supplementary Fig. 7a). By measuring antitermination as a function of perturbed folding pathways, we can then infer the sequence determinants that govern folding transitions. Our strategy was inspired by a comprehensive analysis of glycine riboswitch point mutations^40^ and other approaches that systematically perturb RNA transcripts^41^, and is similar to a recently described in-cell fluorescence-based genetic screen for riboswitch function^42^. Using our mutant libraries, we performed multiplexed *in vitro* transcription in the presence and absence of ZMP and mapped RNA 3’ end distributions by high-throughput sequencing to determine terminator readthrough efficiency (Supplementary Fig. 7a). We find that sequencing measurements approximate those made by gel electrophoresis and are highly reproducible (Supplementary Fig. 7b-e).

To investigate how aptamer/terminator overlap sequence variations can influence ZTP riboswitch folding we constructed a mutagenesis library that varied Y91 and Y94 (Y=U,C) of L3, the respective base pairing partners R29 and R26 in the pseudoknot (R=A,G), and R119 and R114 in the terminator such that the aptamer consensus sequence was preserved^26^ (Fig. 4a). These mutants displayed a range of function, characterized by a difference of the fraction readthrough values in the two conditions (1mM – 0mM ZTP) ranging from 3.7% to 39% (wild-type = 39%) (Supplementary Fig. 8a). As anticipated, mutations that perturb the pseudoknot reduce ZMP-responsiveness and mutations that perturb the terminator increase terminator readthrough (Fig. 4b). The best performing variants contained complete or near-complete Watson-Crick pairing in both the pseudoknot and terminator (Fig. 4c). In contrast, variants with mismatches or wobble pairs typically reduced ZMP-dependent terminator readthrough either by reducing ZMP-responsiveness or increasing background terminator readthrough (Fig. 4c). Among the variants that have Watson-Crick matches within the pseudoknot, the one with the most substantial departure from the WT measurements reversed the native A26:U94/G29:C91 base pairs to G26:C94/A29:U91 and displayed elevated terminator readthrough in the absence and presence of ZMP (Supplementary Fig. 8c-e). Lastly, terminator efficiency in the absence of ZMP was dependent on the position and severity of perturbations; poly-U tract proximal mismatches produced severe termination defects^43^ and strong competing pseudoknot base pairs reduced terminator efficiency Supplementary Fig. 8b).

**Figure 4.**
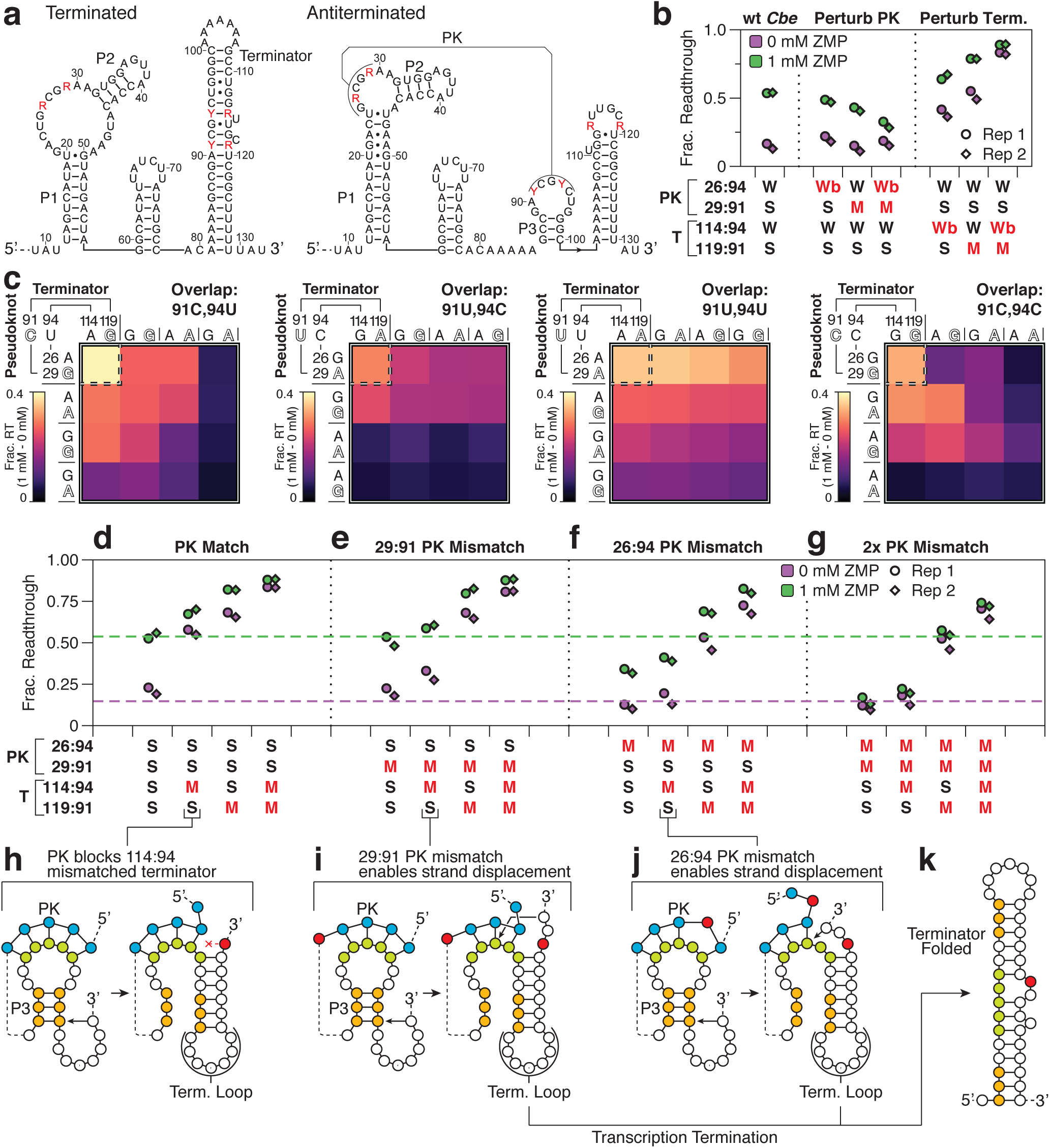
Sequence determinants of efficient terminator formation in the *pfl aptamer pseudoknot*. **(a)** *Cbe* ZTP riboswitch terminated and antiterminated secondary structures depicting the randomization scheme used for aptamer/terminator overlap combinatorial mutagenesis. Randomized nucleotides are shown in red. R=A,G. Y=C,U. **(b)** Plot of fraction readthrough measured from high-throughput sequencing-based transcription assays for pseudoknot mutants as measured in the absence (purple) and presence (green) of 1 mM ZMP. Variant pairing patterns in the pseudoknot (PK) and terminator (T) are annotated as weak (W, A-U), strong (S, G-C), wobble (Wb, G-U), or mismatch (M, A-C). Red annotations indicate deviations from the wild-type pairing pattern. **(c)** Heat maps showing the difference in fraction readthrough between 1 mM and 0 mM ZMP conditions for pseudoknot mutants categorized by the identity of the aptamer/terminator overlap sequence (nts 91 and 94). For each group, the mutant containing perfect Watson-Crick base pairing in both the pseudoknot and terminator are shown by a dashed box (upper left). Competing pseudoknot and terminator base pairs are shown in matching type. **(d-g)** Fraction readthrough for pseudoknot match (d), 29:91 mismatch (e), 26:94 mismatch (f), and 29:91/26:94 mismatch variants (g) with terminator perturbations. Sequences are annotated as in panel (b). Dashed lines indicate observed fraction readthrough for the WT *Cbe* ZTP riboswitch with 0 mM (purple) and 1 mM (green) ZMP. **(h-j)** Models for rescue of a 114:94 terminator mismatch (h) by a 29:91 (i) or 26:94 (j) pseudoknot mismatch. **(k)** Folded terminator secondary structure model. n=2 independent biological replicates are annotated as ‘Rep 1’ and ‘Rep 2’ in panels b and d-g. Heatmaps in panel c are the average of n=2 independent biological replicates. Individual replicate values are compared in Supplementary Figure 7c.

**Figure 4-source data.** Source data for Figure 4 are available in the Northwestern University Arch Institutional Repository (https://doi.org/10.21985/N20F4H).

### *pfl* riboswitch terminator folding requires efficient strand displacement through the pseudoknot

The observed *pfl* riboswitch folding pathway indicates that the pseudoknot folds in the absence of ZMP and is subsequently disrupted during termination. If both P3 and the pseudoknot are folded, the most likely pathway for terminator folding begins with strand displacement through the 3’ side of P3 by the 3’ side of the terminator (Fig. 4h). To test this model, we asked how function of a variant with five GC pseudoknot pairs was altered by mismatches in the pseudoknot, the terminator, or both. In variants with a fully paired pseudoknot, both the poly-U proximal (119A:91C) and poly-U distal (114A:94C) terminator mismatches increased terminator readthrough to >58% in the absence of ZMP (Fig. 4d). While the poly-U proximal terminator mismatch is expected to cause a general termination defect by destabilizing the lower terminator hairpin stem ^40,43^, we reasoned that the poly-U distal mismatch may interfere with terminator folding. If terminator nucleation begins by closing the terminator loop, only one pseudoknot base pair would be disrupted before the poly-U distal mismatch interrupts strand displacement (Fig. 4h), but pseudoknot mismatches might recover terminator function by permitting strand displacement (Fig. 4i-k). In agreement with this model, the 26A:94C and 29A:119C pseudoknot mismatches rescue the poly-U distal terminator mismatch individually (Fig. 4e, f) and in combination (Fig. 4g). Importantly, each individual pseudoknot mutant retains at least partial capacity for ZMP-mediated antitermination, and therefore pseudoknot formation (Fig. 4e, f). Thus, rescue of the poly-U distal terminator mismatch can be linked to recovery of efficient strand displacement through the pseudoknot. In contrast, the inability of individual or double pseudoknot mismatches to rescue terminators with a poly-U proximal mismatch is consistent with a general termination defect (Fig. 4e-g). These data suggest that the apo aptamer forms a stable pseudoknotted fold during transcription that must be broken during termination. Notably, the native *pfl* riboswitch terminator naturally contains bulges that could interfere with displacement of pseudoknot base pairs (Fig. 1b). Deletion of these bulges does not meaningfully impact termination efficiency in the absence of ZMP but reduces terminator readthrough in the presence of ZMP, suggesting that imperfections during strand displacement can tune function (Supplementary Fig. 9).

### Labile P3 base pairing is a critical determinant of termination efficiency in the absence of ZMP

The ZMP-dependence of P1 base pairs that form a ribose zipper with P3 implicates ZMP-mediated P3 stabilization as a determinant of antitermination (Fig. 3). To assess how P3 stability influences termination efficiency, we designed a library that extends P3 by two potential base pairs while preserving terminator base pairing and the ZTP aptamer consensus sequence^26^ (Fig. 5a). The resulting 512 variants assess a model for terminator folding in which nucleation and strand displacement compete with P3 by invading the bottom-most base pairs (Pair 1 and Pair 2) with two invading nucleotides (Invader 1 and Invader 2) to form the top of the terminator stem (Fig. 5b).

**Figure 5.**
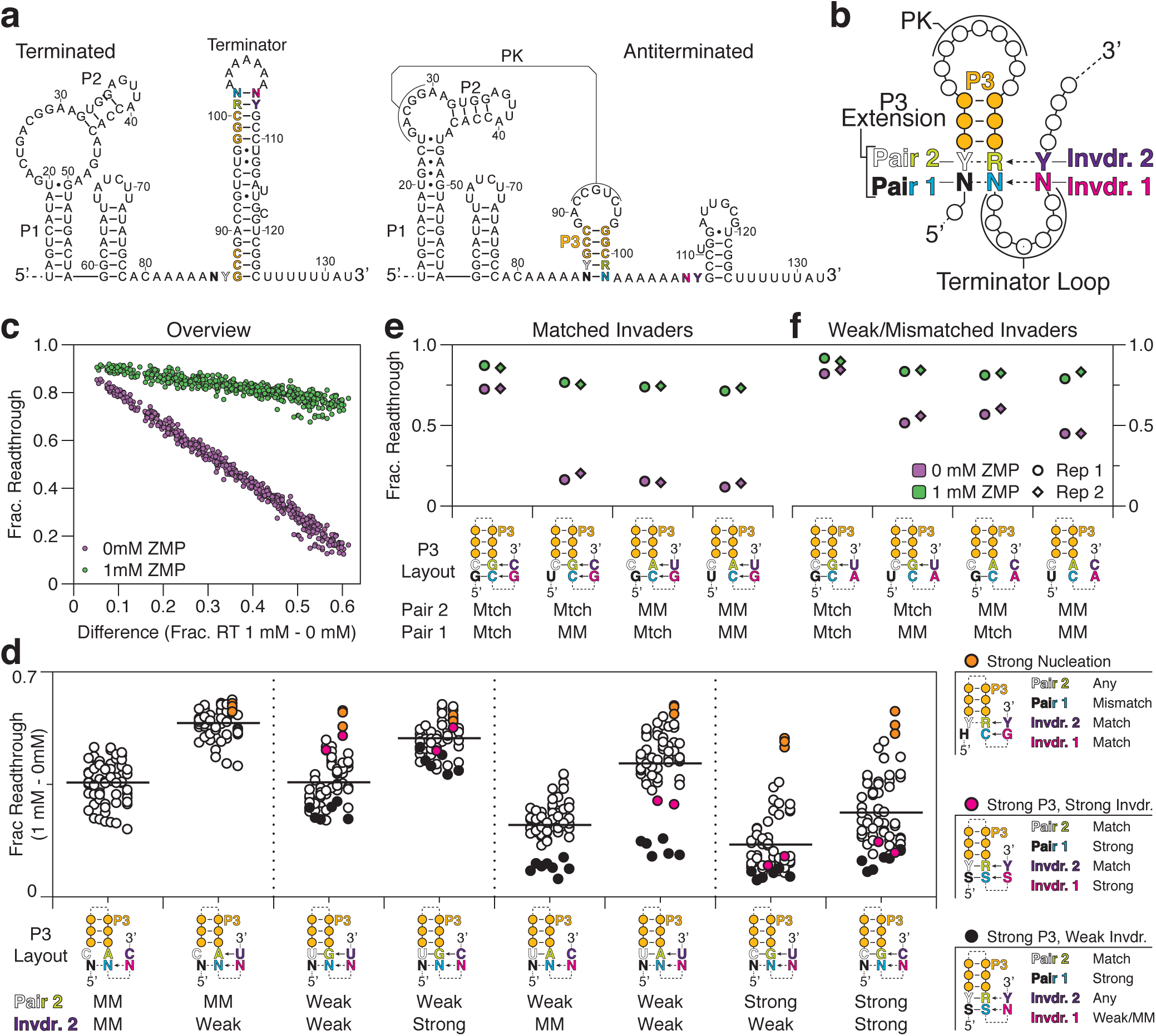
The P3 stem is a critical determinant of ZTP riboswitch termination efficiency. **(a)** *Cbe* ZTP riboswitch terminated and antiterminated secondary structures depicting the randomization scheme used for P3 stem combinatorial mutagenesis. Mutations are color coded according to region. Insertion mutations are not counted in nucleotide numbering to maintain consistent nucleotide numbers. **(b)** Schematic of P3 stem/terminator base pairing competition. Randomized nucleotides are shown using the color codes in (a). Potential base pairs are shown as dashed lines or arrows. Invading nucleotides from the terminator are labeled Invdr. **(c)** Plot of fraction readthrough for P3 mutants as measured in the absence (purple) and presence (green) of 1 mM ZMP ordered by the difference in fraction readthrough observed in 1 mM and 0 mM ZMP conditions. **(d)** Plot of fraction readthrough (1 mM – 0mM) separated by Pair 2 and Invader 2 identity. Solid horizontal lines are the average fraction readthrough (1 mM – 0 mM) of the respective group. P3 Layout for each group is colored as in (b). Notable Pair 2/Invader 2 configurations are plotted in orange (strong nucleation), magenta (strong P3, strong invader), or black (strong P3, weak invader); variable P3 nucleotides are annotated as purine (R, A/G), pyrimidine (Y, U/C), strong (S, G/C), not G (H, A/U/C), or any (N, A/U/G/C). For these configurations, required base pairs are shown with a solid line and optional base pairs are shown in a dashed line. **(e-f)** Plot showing fraction readthrough for select P3 base pairing configurations with matched or weak/mismatched invading nucleotides. P3 Layout for each variant is shown as in (d). n=2 independent biological replicates are annotated as ‘Rep 1’ and ‘Rep 2’ in panels e and f. Panel c and d are the average of n=2 independent biological replicates. Individual replicate values are compared in Supplementary Figure 7d.

**Figure 5 source data.** Source data for Figure 5 are available in the Northwestern University Arch Institutional Repository (https://doi.org/10.21985/N20F4H).

The distribution of all variants ranked by fraction readthrough in the presence and absence of ZMP indicated consistently high terminator readthrough with ZMP, but a range of termination (10-80% terminator readthrough) in the absence of ZMP (Fig. 5c). Thus, for this mutant group, ZMP-dependent activation is defined by termination efficiency. Classification of P3 variants by the strength of Pair 2, which can directly extend P3 by up to 1 bp, reveals that improved base pairing in P3 tends to reduce terminator efficiency (Fig. 5d). Furthermore, for each Pair 2 variant, the presence of a complementary Invader 2 nucleotide improves termination efficiency in the absence of ZMP, and thus riboswitch function, relative to a mismatched Invader 2 (Fig. 5d). Interestingly, we also observed that sequences in which a G in Invader 1 can pair with a 3’ C in Pair 1 always results in an optimally functioning variant within sequence groups, but not when the G-C pair orientation is reversed (Fig. 5d orange points, Supplementary Fig. 10d). A closer analysis of the interplay between several P3 base pairing configurations and perfectly matched invading nucleotides yield highly functional riboswitches except when extension of P3 by two G-C pairs make it inaccessible to strand displacement (Fig. 5e). Weak or mismatched invading pairs reduce termination efficiency in the absence of ZMP and remain sensitive to strong P3 base pairing (Fig. 5f). Overall, these data uncover sequence features of efficient terminator nucleation in the absence of ZMP and identify the accessibility of P3 to strand displacement as a critical determinant of termination efficiency.

### *pfl* aptamer mutants are prone to misfolding

In a mutagenesis experiment designed to perturb pseudoknot-contacting nucleotides, we randomized A31 and A40 (Fig. 3b) alongside A90, which was observed to form a base triple with a pseudoknot pair in the *Thermosinus carboxydivorans pfl* riboswitch structure ^30^ (Supplementary Fig. 11a-c). To complement N90 mutations in the terminator hairpin we randomized U120 (Supplementary Fig. 11b). Surprisingly, all non-WT 90:120 pairs increased terminator readthrough in the absence of ZMP and 90Y:120R pairs functioned as poorly as most mismatches (Supplementary Fig. 11d). We identified three possible causes for these defects: First, minimum free energy models of the local P3/L3 sequence suggest that nt 90 G, U, and C variants can form favorable off-pathway hairpins that compete with P3 and could interfere with terminator nucleation (Supplementary Fig. 11e). Second, the 90U variant may form an alternate pseudoknot or extend the natural pseudoknot by one base pair (Supplementary Fig. 11f). In agreement with this interpretation, three mutations at 31 and 40 that are predicted to occlude the pseudoknot by causing a P2 misfold (Supplementary Fig. 11G, i-k) partially restore termination efficiency (Supplementary Fig. 11d, 90U:120A red points). Third, the mutations that partially rescue 90U:120A variants have no effect on 90U:120G variants, suggesting that the 120G mutation causes a terminator defect, though its origin is unclear (Supplementary Fig. 11d, 90U:120G red points). Lastly, the 31U mutation causes elevated terminator readthrough which may be related to its capacity to extend P2 by two base pairs, thereby distorting J1/2 and J2/3 (Supplementary Fig. 11h, i). The tendency of these relatively minor perturbations to profoundly impact *pfl* riboswitch function may be related to the substantial challenge of folding and associating two separate aptamer domains in time for ZMP recognition during a short window of transcription (Figs. 2e, 7).

### Context-dependent sequence composition guides proper P3 folding

Our structural and functional analyses support a central role for P3 in ZTP riboswitch antitermination. Notably, 14 of 15 nucleotides that comprise P3/L3 are conserved in identity while the remaining nucleotide (A90 of *Cbe pfl*), is present in >97% of ZTP aptamers but without nucleotide preference^26^ and was observed in three distinct conformations^28–30^. Nonetheless, mutation of this position is associated with profound terminator defects (Supplementary Fig. 11c, d), some of which may be due to competing P3 misfolds (Fig. 6a-c). To ask whether the nucleotide context in P3/L3 could govern a conservation pattern that avoids off-pathway folding, we assessed the prevalence of an eight nucleotide ‘misfold motif’ that is predicted to form an off-pathway fold if the second L3 nucleotide (L3n2) is C or, to a lesser extent, U (Fig. 6d). To do this, we binned 532 ZTP riboswitch sequences from fully sequenced bacterial genomes^26^ by the presence of the seven nucleotides that yield the misfold motif if L3n2 is a pyrimidine (Fig. 6d) and the number of contiguous P3 pairs as a proxy for the favorability of the correct P3 fold, and then determined the nucleotide distribution at L3n2. In the aggregate set of ZTP riboswitch sequences A, U, and C were present at L3n2 at approximately equal frequencies. However, among riboswitches that would complete the misfold motif if L3n2=Y, >91 % of sequences with a 3 bp P3 stem contain an A at L3n2 (Fig. 6e). Based on our analysis above, this suggests the sequence at this position is constrained to prevent possible off pathway folds. Interestingly, this constraint is relaxed as P3 grows in length - the frequency of A at L3n2 was reduced to >79% for sequences with a 4 bp P3 stem and sequences with a 5 bp P3 stem favor L3n2 pyrimidines comparably to sequences without any capacity for the misfold motif (Fig. 6e). This suggests that the sequence constraints that avoid the potential for misfolding with a weaker P3 stem may not be necessary if P3 is highly favorable. Taken together, these observations suggest that in specific sequence and structure contexts there is a selective pressure within L3 to prevent misfolding of a crucial riboswitch structure.

**Figure 6.**
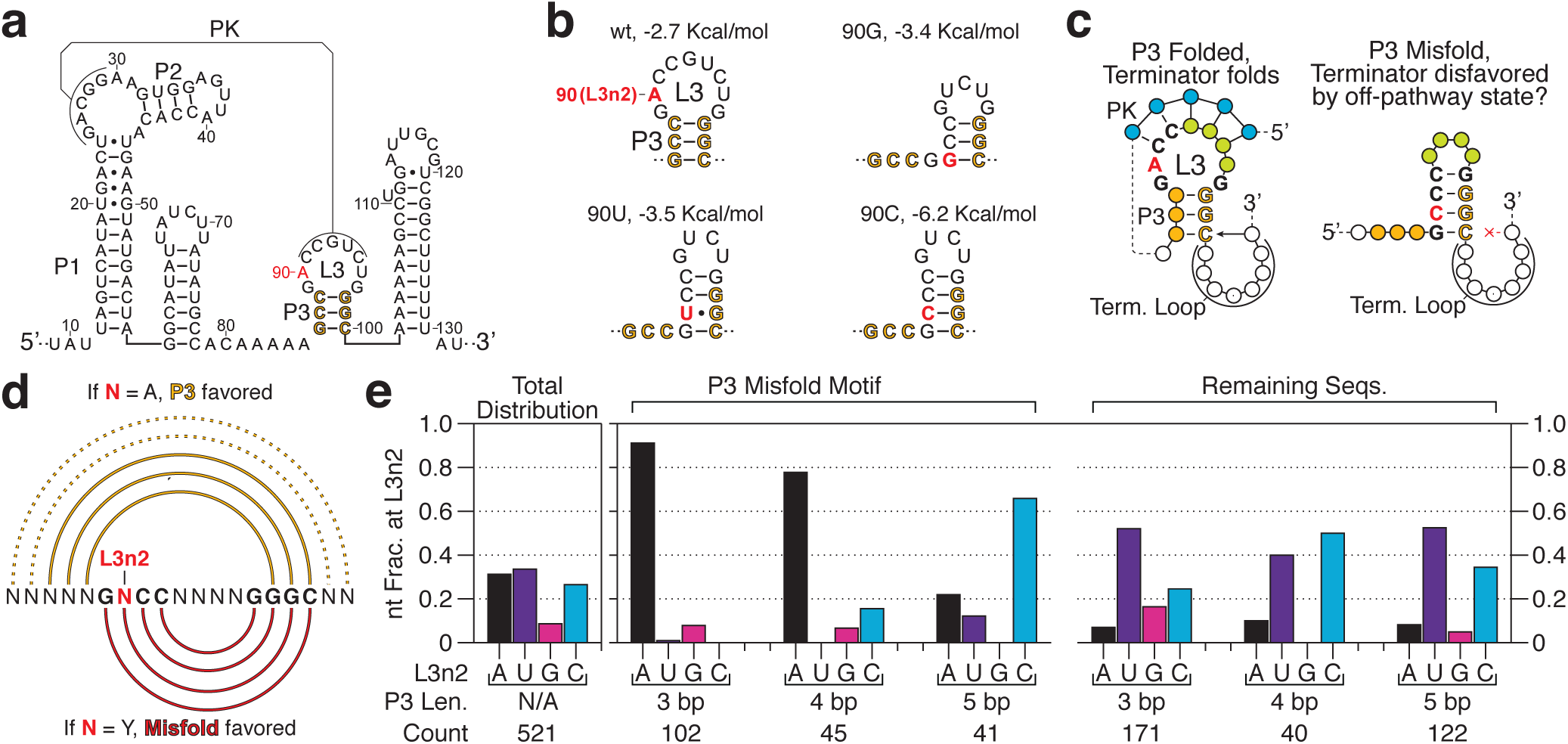
Context-dependent nucleotide frequency in P3 sequences suggests selective pressures for on-pathway folding. **(a)** *Cbe* ZTP riboswitch antiterminated secondary structure. P3 stem nucleotides are gold and A90 is red. **(b)** Structure predictions for each N90 variant of the P3 region (nts 86-100) with an associated ΔG predicted by the RNAstructure Fold command. **(c)** A model for how a P3 misfold could interfere with terminator folding. Native P3 base pairs are indicated by gold coloring; The sequence region comprising the misfold motif is shown in bold text with a defined sequence, the red nucleotide designates L3 nucleotide 2 (L3n2) and corresponds to A90 in the WT *Cbe pfl* ZTP aptamer. **(d)** P3 folding in the context of the P3 misfold motif sequence. The sequence region comprising the misfold motif is shown in bold text with a defined sequence except for the red ‘N’, which designates L3 nucleotide 2 (L3n2) and corresponds to A90 in the wt *Cbe pfl* ZTP aptamer. A complete match to defined nucleotides designates a sequence as having the capacity to misfold if L3n2=Y. All nucleotides that were not specified when classifying P3/L3 sequences are labeled ‘N’. The native P3 fold is shown above the sequence text in gold, where arcs between two bases indicate pairing: solid lines designate critical P3 base pairs and dashed lines designate optional P3 base pairs observed in some ZTP riboswitch sequences ^26^. The red lines below the sequence text illustrate the modeled misfold structure that is favored over the native P3 structure when N=Y. **(e)** Nucleotide frequency at position 90 for native ZTP riboswitch sequences in aggregate and binned by the presence of the ‘misfold motif’ defined in (d) and the length of the P3 stem. Sequences containing non-conserved insertions or a maximum contiguous P3 stem of less than three base pairs (11 of 532 sequences) were not considered.

**Figure 6-source data.** Source data for Figure 6 are available in the Northwestern University Arch Institutional Repository (https://doi.org/10.21985/N20F4H).

## Discussion

Here we have shown how intermediate folds of the *Cbe pfl* ZTP riboswitch mediate its function as a ligand-dependent transcription antiterminator. Nascent RNA structure probing revealed a positional roadmap of ZTP riboswitch folds comprising the formation of a transient hairpin, followed by aptamer organization, ligand-mediated aptamer stabilization, and bifurcation into terminated or antiterminated states (Fig. 7). We found that the *pfl* aptamer pseudoknot folds independently of ZMP during transcription, but that ZMP binding stabilizes non-Watson-Crick P1 base pairs that directly contact P3. We then used a combinatorial mutagenesis approach to reveal that the *pfl* ZTP riboswitch terminator hairpin nucleates by invading P3 and propagates by strand displacement through pseudoknot base pairs. Together, these analyses uncover sequence determinants that govern *pfl* ZTP riboswitch terminator folding in the absence of ZMP and suggest a mechanism by which ZMP binding converts P3 into a rigid barrier to terminator nucleation.

**Figure 7.**
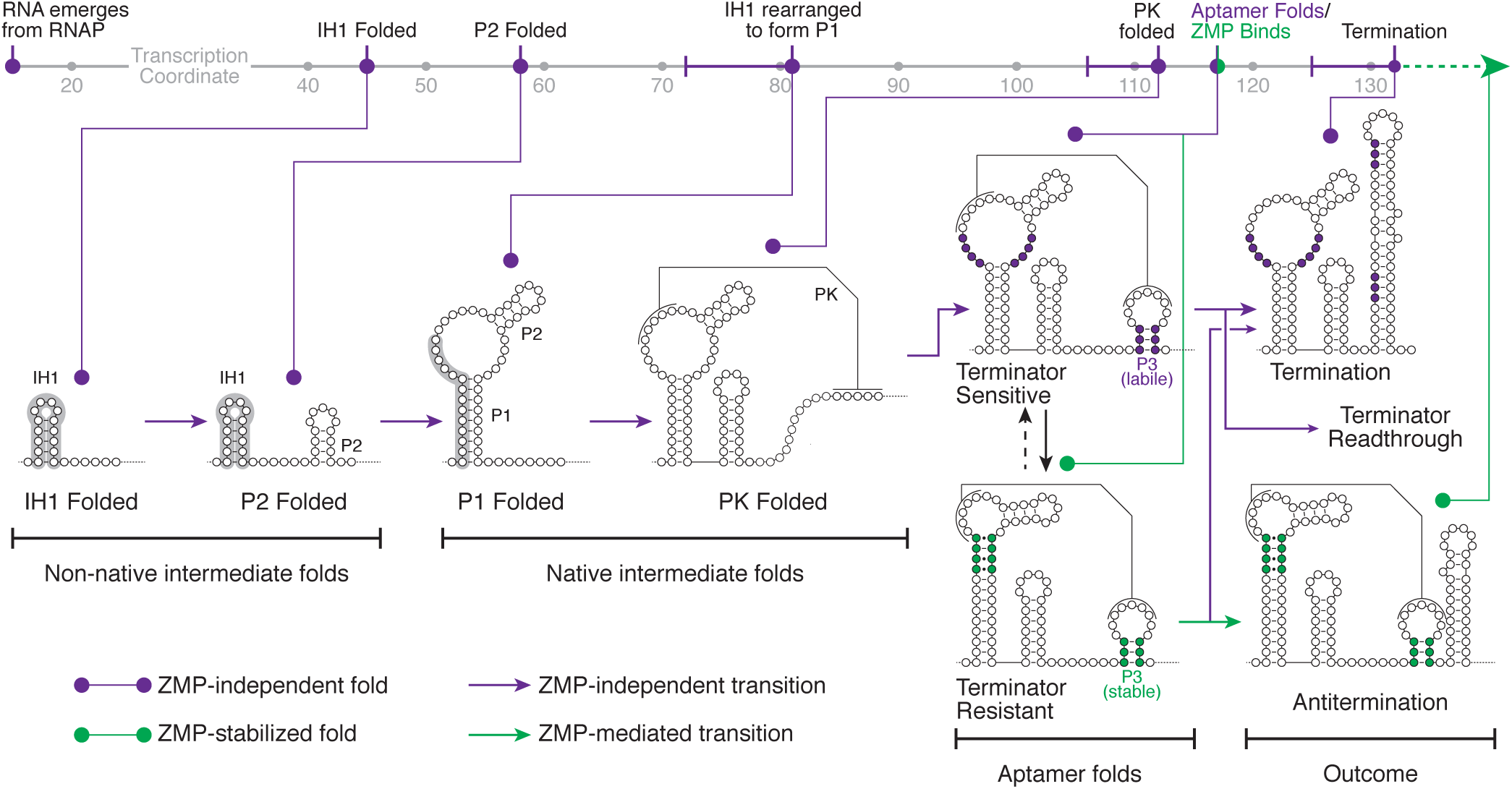
A model for ZTP riboswitch folding. ZTP riboswitch folding intermediates as observed by cotranscriptional SHAPE-Seq are plotted by transcription coordinate. Purple indicates ZMP-independent folds and transitions; Green indicates ZMP-stabilized folds or ZMP-mediated transitions. Initial ZTP aptamer folding comprises the formation of a transient intermediate hairpin (IH1) that rearranges to form P1. Upon folding of both the pseudoknot (PK) and P3, the ZTP aptamer is competent to bind ZMP. In the absence of ZMP, terminator nucleation sequentially disrupts P3 and PK to fold the terminator hairpin and terminate transcription. In the presence of ZMP, the aptamer can become trapped in a stable fold that renders the P3 stem resistant to terminator nucleation, thereby driving antitermination.

Our analysis of the Cbe *pfl* ZTP riboswitch revealed that terminator folding proceeds by strand displacement through the ZTP aptamer. Terminator folding is efficient if the pairing interactions in the pseudoknot and P3 stem are balanced with terminator base pairing. Nucleotide compositions that favor either the aptamer or the terminator structure tip this balance and lead to non-functional switches (Figs. 4, 5). These features of *pfl* riboswitch folding are linked to its capacity for stable pseudoknot formation in the absence of ZMP and are similar to observations that the *crcB* fluoride aptamer adopts a psuedoknotted fold that must be disrupted during termination ^22,23^. In contrast, the *Faecalibacterium prausnitzii* class III preQ_1_ riboswitch was found to dynamically sequester its ribosome binding site by rapid docking and undocking within a pseudoknot ^44^, suggesting that distinct regulatory contexts require specific folding dynamics. In the case of the *pfl* ZTP and *crcB* fluoride aptamers, stable pseudoknot formation may be important for ligand recognition within the short window of transcription, but consequently requires efficient strand displacement for terminator folding.

Given that the *pfl* riboswitch terminator has evolved to efficiently displace a pseudoknotted hairpin, how does ZMP binding favor antitermination? First, ZMP-recognition was shown to join P3 and the pseudoknot in a continuous helical stack by forming a Hoogsteen edge-Watson-Crick edge pair with a highly conserved U in L3^28–30^. In addition, in-line probing measurements^26^ and nascent RNA structure probing (Fig. 3) show that several ZMP-responsive nucleotides localize to a non-Watson-Crick extension of P1 that forms a ribose zipper with P3^28–30^. Thus, ZMP binding could mediate antitermination both by extending the P3 helical stack^28–30^ and by facilitating intersubdomain contacts between P1 and P3. Notably, the former is similar to a mechanism proposed for adenine-mediated stabilization of the pbuE aptamer P1 stem against terminator strand displacement^18^. Consistent with the localization of ZMP-stabilized interactions to P3, *pfl* riboswitch insertion mutants that vary P3 and invading terminator nucleotides are exquisitely sensitive to the competition between P3 base pairs and terminator nucleation (Fig. 5). Furthermore, we observed that an otherwise stable P3 stem can be efficiently disrupted if terminator folding can be nucleated by the formation of a base pair at the 3’ base of the stem. This is similar to ‘toehold’ strand displacement in synthetic DNA hybridization systems where efficient DNA strand displacement is seeded through interactions in unpaired DNA regions^45,46^. While this mode for nucleating strand displacement is local relative to classical ‘toeholds’, the underlying principle was also recently reported for the *pbuE* adenine riboswitch^47^ and appear to be general. Interestingly, ZMP binding can still antiterminate transcription in the context of variants with the capacity for efficient terminator nucleation (Fig. 5). This suggests that ZMP-dependent stabilization of P3 is versatile and can block both terminator nucleation and propagation of strand displacement. Overall these considerations between balancing the two mutually exclusive riboswitch folds and including motifs that allow folding pathways to traverse barriers caused by imbalances could constrain riboswitch sequence evolution and greatly inform riboswitch design.

An important constraint to ZTP-mediated antitermination is the narrow window of transcription coordinates in which the aptamer can bind ZMP before terminator folding. The *pfl* aptamer is competent to bind ZMP ~15 nts before RNAP reaches the primary termination site, but only ~7 nucleotides before terminator nucleation can begin to compete with the aptamer fold (Fig. 2). ZMP binding within this 15 nt window could be promoted by several mechanisms. First, pseudoknot base pairs can form up to 11 nucleotide addition cycles before ZMP can bind. Second, transcription pausing^48,49^ has been implicated in riboswitch folding and ligand recognition, and an appropriately positioned pause could extend the time for ZMP binding^21,25^. However, rigorous evaluation of pausing would require use of the cognate RNAP^50^. Finally, the observation that antitermination is efficient given sufficient ZMP suggests that ZTP aptamer folding is frequently successful and inefficiency in ligand binding may be tolerated over many rounds of transcription.

Importantly, our findings only apply to a cotranscriptional folding regime; equilibrium refolding favors terminator formation even in the presence of ZMP. Thus, the order of folding is essential for function because antitermination depends on cotranscriptional entry into a kinetically trapped holo aptamer fold. In this regard, the ZTP riboswitch behaves similarly to the *Bacillus cereus crcB* fluoride^22^ and *Escherichia coli thiC* thiamine pyrophosphate (TPP) riboswitches^25^. Notably, the apo *pfl* aptamer requires only a single additional A:U or G:C pair to trap the riboswitch in an antiterminated state without ligand recognition. This is consistent with an analysis of the *Clostridium tetani* glycine riboswitch type-1 singlet which concluded that meaningful ligand-dependent changes in gene expression require finely tuned expression platform helices that are close in energy^40^. Interestingly, in the case of the *pfl* riboswitch, sequences that promote highly efficient terminator nucleation can circumvent an artificially stabilized aptamer. Thus, the *pfl* riboswitch folding pathway inherently assumes an ON decision and relies on both the thermodynamic favorability of the terminator structure, and the kinetic efficiency of strand displacement, to reject this structural assumption in the absence of ligand.

Our analysis of the *Cbe pfl* riboswitch provides evidence for two proposed characteristics of cotranscriptional RNA folding pathways: the presence of temporary helices and the avoidance of competitor helices^51^. The formation of local transient folds is unexpected for transcriptional riboswitches because function requires successful aptamer folding. Nonetheless, the temporary helix IH1 precedes *pfl* aptamer folding and is one of the few experimentally observed transient intermediate folds^7,22,52–54^ (Fig. 2). Interestingly, the IH1 structure is not explicitly encoded in the *pfl* aptamer consensus sequence but is enriched for by the high GC content of J1/2 and the separation of the sequences that comprise P1 by ~30 bp. The simplicity of these characteristics suggests that IH1-like structures may be prevalent in other non-coding RNAs. In addition to its capacity for transient folds, the ZTP aptamer exhibits a context-dependent sequence preference that may avoid a misfolded alternative structure to P3 that would preclude ZMP recognition (Fig. 6). Importantly, sequence preference at this position is dependent on both the potential for misfolding given the local sequence context and the favorability of the native P3 structure. While these findings are limited in scope, they support a general principle that RNA sequences are selected both for their functional structure and for efficient folding pathways to guide formation of that structure ^3^.

## Materials and Methods

### Standard DNA template preparation

Linear DNA template for *in vitro* transcription were prepared by PCR amplification as previously described^55^. Briefly, five 100 μl reactions containing 82.25 μl of water, 10 μl Thermo Pol Buffer (New England Biolabs, Ipswich, MA), 1.25 μl of 10 mM dNTPs (New England Biolabs), 2.5 μl of 10 μM oligonucleotide A (forward primer; Supplementary Table 4), 2.5 μl of 10 μM oligonucleotide B (reverse primer; Supplementary Table 4), 1 μl of Vent Exo-DNA polymerase (New England Biolabs), and 0.5 μl of plasmid DNA (Supplementary Table 3) were subjected to 30 cycles of PCR. Terminal biotin roadblock DNA templates were prepared as above but using either oligonucleotide D or oligonucleotide E (Supplementary Table 4) as the reverse primer. Reactions were pooled and precipitated by adding 50 μl of 3M sodium acetate (NaOAc) pH 5.5 and 1 mL of cold 100% ethanol (EtOH) and incubating at − 80C for 15 min. After centrifugation, the precipitated pellet was washed once with 1.5 mL 70% EtOH (v/v), dried using a SpeedVac, and dissolved in 30 μl of water. The template was run on a 1% agarose gel and extracted using a QIAquick Gel Extraction kit (Qiagen, Hilden, Germany). DNA template concentration was determined by a Qubit 3.0 Fluorometer (Life Technologies, Carlsbad, CA).

### Biotinylated DNA template preparation

Linear DNA templates containing approximately 1 internal biotin modification were prepared by PCR amplification and gel extraction as previously described^34^. Briefly, PCR amplification was performed as described above except that in place of a standard dNTP mixture, each dNTP and corresponding biotin-11-dNTP was added to a total of 100 nmol such that ~1 biotin-11-dNTP is incorporated with in the transcribed region of each DNA duplex as described previously^34^. Biotin-11-dATP and biotin-11-dGTP were purchased from Perkin Elmer (Waltham, MA). Biotin-11-dCTP and biotin-11-dUTP were purchased from Biotium (Fremont, CA).

### Combinatorial mutagenesis DNA template preparation

Linear DNA templates for combinatorial mutagenesis experiments were prepared by PCR amplification of gel-purified ‘Ultramer’ oligonucleotides (Integrated DNA technologies, Coralville, IA) (Supplementary Table 5). Ultramer oligonucleotides were first converted to double-stranded DNA in a 100 μl PCR containing 84.5 μl water, 10 μl 10X ThermoPol Buffer, 2 μl 10 mM dNTPs, 0.25 μl of 100 μM oligo F (forward primer; Supplementary Table 4), 0.25 μl of 100 μM oligo E (reverse primer; Supplementary Table 4), and 2 μl of Vent Exo-DNA polymerase. Thermal cycling was performed for 8 cycles. The primers in this reaction appended additional sequence to the ultramer. The PCR was purified using a QIAquick PCR purification kit (Qiagen) and quantified using a Qubit 3.0 Fluorometer. A second preparatory amplification was then performed using oligonucleotides G and E (Supplementary Table 4) followed by precipitation and gel extraction as described above for Standard DNA Templates. The resulting product was quantified using a Qubit 3.0 Fluorometer.

### *in vitro* transcription (without roadblocking)

Open promoter complexes were formed as previously described^34^ by incubating 100 nM DNA template and 2 U of *E. coli* RNAP holoenzyme (New England Biolabs) in transcription buffer (20 mM tris(hydroxymethyl)aminomethane hydrochloride (Tris-HCl) pH 8.0, 0.1 mM ethylenediaminetetraacetic acid (EDTA), 1 mM dithiothreitol (DTT) and 50 mM potassium chloride (KCl)), 0.2 mg/ml bovine serum albumin (BSA), and 500 μM High Purity ATP, GTP, CTP and UTP (GE Life Sciences) for 10 min at 37C. When present, 5-aminoimidazole-4-carboxamide-1-β-D-ribofuranosyl 5’-monophosphate (ZMP; Sigma Aldrich, St. Louis, MO) at a stock concentration of 50 mM in dimethyl sulfoxide (DMSO) was added to a final concentration of 1 mM ZMP and 2% (v/v) DMSO. When ZMP was not included DMSO was added to 2% (v/v). Single-round transcription was initiated upon addition of MgCl_2_ to 10 mM and rifampicin (Sigma Aldrich) to 10 μg/ml and a total reaction volume of 25 μl. Transcription proceeded for 30 s before 75 μl of TRIzol solution (Life Technologies) was added and RNAs were extracted according to the manufacturer’s protocol.

### *in vitro* transcription with random streptavidin roadblocks and chemical probing (cotranscriptional SHAPE-Seq)

*in vitro* transcription for cotransctiptional SHAPE-Seq was performed as described^34^ for single-length DNA templates except that reactions were scaled to 50 μl and the open promoter complex protocol was modified to accommodate streptavidin binding as previously described^34^. Briefly, after incubation of reactions at 37C for 7.5 min to form open promoter complexes, SAv monomer (Promega, Fitchburg, WI) was then added to 2.5 μM and reactions were incubated for an additional 7.5 min before transcription was initiated by adding MgCl_2_ to 10 mM and rifampicin (Sigma Aldrich) to 10 μg/ml for total reaction volume of 50 μl. Transcription proceeded for 30 s to permit stable distribution of transcription elongation complexes before reactions were split into 25 μl aliquots and mixed with 2.79 μl of 400 mM Benzoyl Cyanide (BzCN; Pfaltz & Bauer, Waterbury, CT) dissolved in anhydrous DMSO ((+) sample) or mixed with anhydrous DMSO ((−) sample) for ~2 s before transcription was stopped by the addition of 75 μl TRIzol solution (Life Technologies) and RNAs were extracted according to the manufacturer’s protocol. DNA template was degraded by incubation in 20 μl 1x DNase I buffer (New England Biolabs) containing 1 U DNase I (New England Biolabs) at 37C for 30 min. 30 μl of water and 150 μl TRIzol were added to stop the reaction and RNAs were extracted according to the manufacturer’s protocol. The resulting RNA was dissolved in 10 μl of 10% DMSO.

*in vitro* transcription for equilibrium refolding experiments were performed as above, except that all reactions contained ZMP to promote stable distribution of elongation complexes across all positions, transcription was stopped without performing chemical probing, and the resulting RNAs were resuspended in 25 μl water. Under this purification protocol the ZMP included during initial RNA synthesis should be completely depleted during the two subsequent phased extractions, as is evidenced by the difference between the equilibrium refolded and cotranscriptionally-folded matrices (compare Fig. 2a and Supplementary Fig. 1a). Equilibrium refolding was performed by denaturing at 95C for 2 min, snap cooling on ice for 1 min, and adding transcription buffer, 500 μM NTPs, 10 mM MgCl_2_, and either ZMP to 1mM ZMP/2% DMSO or 2% DMSO before incubation at 37C for 20 min. SHAPE modification with BzCN was performed as described above before addition of 30 μl water and 150 μl TRIzol, extraction according to the manufacturer’s protocol and resuspension in 10 μl of 10% DMSO.

### *in vitro* transcription with a terminal roadblock (combinatorial mutagenesis)

We chose to include a terminal biotin-streptavidin roadblock as part of our combinatorial mutagenesis protocol so that terminator run-through RNAs would appear as cluster of 3’ ends ~10 nts upstream of the DNA template end. After incubating reactions at 37C for 7.5 min to form open promoter complexes, SAv monomer (Promega) was then added to 2.5 μM and reactions were incubated for an additional 7.5 min before transcription was initiated by adding MgCl_2_ to 10 mM and rifampicin (Sigma Aldrich) to 10 μg/ml for total reaction volume of 25 μl. Transcription proceeded for 30 s before 75 μl of TRIzol solution (Life Technologies) was added and RNAs were extracted according to the manufacturer’s protocol, DNaseI treated, TRIzol extracted a second time as described above.

### Gel Electrophoresis and Analysis of Gel Images

Extracted RNAs were fractionated by denaturing 7.0 M urea polyacrylamide gel electrophoresis (10 or 12% polyacrylamide). Gels were stained with SYBR Gold (Life Technologies), imaged with a Bio-Rad ChemiDoc Touch Imaging System, and quantified with Image Lab (Bio-Rad, Hercules, CA). Fraction readthrough was determined by dividing the intensity of the terminator readthrough band by sum of the terminator readthrough and terminated bands.

### Sequencing library preparation

Sequencing libraries for cotranscriptional SHAPE-seq were prepared either as previously described^56^ or with a modified protocol that uses Superscript IV (SSIV) for reverse transcription. All combinatorial mutagenesis libraries were prepared using the modified SSIV protocol. For convenience, all protocol modifications are described below in the context of the complete original protocol.

### RNA 3’ linker adenylation and ligation

5’-Phosphorylated linker (Oligonucleotide K, Supplementary Table 6) was adenylated using a 5’ DNA Adenylation Kit (New England Biolabs) at 20x scale and purified by TRIzol extraction as previously described^56^. RNA 3’ ligation was performed by first combining 10 μl extracted RNAs in 10% DMSO with 0.5 μl of SuperaseIN (Life Technologies), 6 μl 50% PEG 8000, 2 μl of 10X T4 RNA Ligase Buffer (New England Biolabs), 1 μl of 2 μM 5’-adenylated RNA linker, and 0.5 μl of T4 RNA ligase, truncated KQ (New England Biolabs) and mixing by pipetting. 0.5 μl of T4 RNA ligase 2, truncated KQ (New England Biolabs) was then added and the reaction was mixed by pipetting again and incubated at 25C for 3 hrs.

### Reverse Transcription

Following linker ligation, RNAs were precipitated by adding 130 μl RNase-free water, 15 μl 3M NaOAc pH 5.5, 1 μl 20 mg/ml glycogen, and 450 μl of 100% EtOH and storing at −80C for 30 min. Following centrifugation, pellets were washed once with 500 μl 70% EtOH (v/v), residual ethanol was removed. For Superscript III reverse transcription, pellets were resuspended in 10 ul RNase-free water and 3 μl of reverse transcription primer (Oligonucleotide L, Supplementary Table 6) was added. Samples were then denatured at 95C for 2 min, incubated at 65C for 5 min, briefly centrifuged to pull all liquid to the bottom of the tube, and placed on ice. 7 μl of Superscript III reverse transcription master mix (containing 4 μl of 5x First Strand Buffer (Life Technologies), 1 μl of 100 mM DTT, 1 μl 10 mM dNTPs, 0.5 μl RNase OUT (Invitrogen, Waltham, MA), and 0.5 μl Superscript III) was added to each sample and samples were mixed before being placed at 45C. Reactions were incubated at 45C for 1 min, 52C for 25 min, and 65C for 5 min to deactivate Superscript III. For Superscript IV reverse transcription, RNAs were precipitated as described for Superscript III but resuspended in 9.5 μl of RNase-free water before 3 μl of reverse transcription primer (Oligonucleotide L, Supplementary Table 6) was added. Samples were then denatured at 95C for 2 min, incubated at 65C for 5 min, briefly centrifuged to pull all liquid to the bottom of the tube, and placed on ice. 7.5 μl of Superscript IV reverse transcription master mix (containing 4 μl of 5x SSIV Buffer (Life Technologies), 1 μl of 100 mM DTT, 1 μl 10 mM dNTPs, 0.5 μl RNase OUT (Invitrogen), and 1 μl Superscript IV) was added to each sample and samples were mixed before being placed at 45C. Reactions were incubated at 45C for 1 min, 52C for 25 min, and 65C for 5 min, and 80C for 10 min to deactivate Superscript IV. Following heat inactivation of Superscript III or Superscript IV, 1 μl 4M sodium hydroxide (NaOH) was added and samples were heated at 95C for 5 min to hydrolyze RNA. Samples were then partially neutralized by the addition of 2 μl 1M hydrochloric acid (HCl) and precipitated by adding 69 μl 100% EtOH and storing at −80C for 15 min. Samples were centrifuged for 15 min at 4C and pellets were washed by adding 500 μl of 70% EtOH and inverting the tubes several times. After removing residual ethanol pellets were dissolved in 22.5 μl of RNase-free water.

### Adapter ligation

Ligation of high-throughput sequencing adapter was performed as previously described^56^. Briefly, the dissolved cDNA was mixed with 3 μl of 10x CircLigase Buffer (Epicentre), 1.5 μl of 50 mM MnCl_2_, 1.5 μl of 1 mM ATP 0.5 μl of 100 μM DNA adapter (Oligonucleotide M, Supplementary Table 6), and 1 μl of CircLigase I (Epicentre, Madison, WI). Ligation reactions were incubated at 60C for 2 h and the ligase was heat inactivated by incubating at 80C for 10 min. DNA was precipitated by adding 70 μl nuclease-free water, 10 μl 3M NaOAc pH 5.5, 1 μl 20 mg/ml glycogen, and 300 μl of 100% EtOH and storing at −80C for 30 min before centrifugation. The pellets were dissolved in 20 μl of nuclease-free water, purified using 36 μl of Agencourt XP beads (Beckman Coulter, Brea, CA) according to the manufacturer’s protocol, and eluted with 20 μl of 1X TE buffer.

### Quality Analysis

Quality analysis for cotranscriptional SHAPE-Seq was performed as previously described^56^ by generating fluorescently labeled dsDNA libraries using oligonucleotides O, P, Q and R or S (Supplementary Table 6). Samples were analyzed by capillary electrophoresis using an ABI 3730xl DNA Analyzer. Combinatorial mutagenesis libraries were also assessed for quality in this way.

### Sequencing Library Preparation

Sequencing library preparation for cotranscriptional SHAPE-Seq was performed as previously described^56^. Briefly, 3 μl of ssDNA library (+) and (−) channels was separately mixed with 33.5 μl of nuclease-free water, 10 μl 5x phusion Buffer (New England BioLabs), 0.5 μl of 10 mM dNTPs, 0.25 μl of 100 μM TruSeq indexing primer (Oligonucleotide T, Supplementary Table 6), 2 μl of 0.1 μM of channel-specific selection primer (Oligonucleotides R and S, Supplementary Table 6), and 0.5 μl of Phusion DNA polymerase (New England BioLabs). Amplification was performed with an annealing temperature of 65C and an extension time of 15 s. After 15 cycles, 0.25 μl of primer PE_F (Oligonucleotide Q, Supplementary Table 6) was added and libraries were amplified for an additional 10 cycles. Following amplification, libraries were allowed to cool to 4C completely before the addition of 0.25 μl of ExoI (New England Biolabs) and incubation at 37C to degrade excess oligonucleotides. ExoI was heat inactivated by incubating at 80C for 20 min. Libraries were then mixed with 90 μl of Agencourt XP beads (Beckman Coulter), purified according to the manufacturer’s protocol and eluted in 20 μl of 1X TE buffer. The resulting sequencing libraries were quantified using a Qubit 3.0 Fluorometer (Life Technologies) and molarity was estimated using the length distribution observed in Quality Analysis. Sequencing libraries for combinatorial mutagenesis was performed as described above except that (+)/(−) channel barcoding was arbitrarily assigned because each library was given a unique TruSeq barcode.

### Cotranscriptional SHAPE-Seq Sequencing and analysis

Sequencing of cotranscriptional SHAPE-Seq libraries was performed by the NUSeq Core on an Illumina NextSeq500 using either 2×36 or 2×37 bp paired end reads with 30% PhiX. Cotranscriptional SHAPE-Seq data analysis was performed using Spats v1.0.1 as previously described^22^ except that one mismatch was permitted during alignment following the observation that truncations at several nucleotides were enriched for a terminal mutation.

### Combinatorial Mutagenesis Data Sequencing and Analysis

Sequencing of combinatorial mutagenesis libraries was performed on an Illumina MiSeq using a MiSeq Reagent Kit v3 (150-cycle). Libraries were loaded with a density of approximately 1000 K/mm^2^ and sequenced with a cycle configuration of either Read1:37, Index:6, Read2:132 or Read1:35, Index:6, Read2:134 and included 10% PhiX. Asymmetrical reads were used so that nearly the entire riboswitch was included in read 2 and the transcript end was contained in read 1. Alignment was performed using custom software available at https://github.com/LucksLab/LucksLab_Publications/tree/master/Strobel_ZTP_Riboswitch. For mutants without insertions, reads were required to contain a perfect target match between nucleotides 11 and 130 of the riboswitch. Alignment to the riboswitch leader (nts 1-10) was not required because elevated Superscript IV dropoff in this A/U-rich region reduced the number of alignments, and omission of reads lacking this sequence recovered these reads but did not impact measurements (Supplementary Fig. 7f, g). Control analyses of non-insertion mutants that omitted the poly-uridine tract from alignment did not impact measurements (Supplementary Fig. 7f, g), therefore insertion mutants were only required to align through two poly-uridine tract nucleotides to permit usage of 150-cycle v3 MiSeq Reagent Kits. For all mutants, Single mismatches were permitted in Read 1 beyond nt 130 provided that the read could be unambiguously identified as terminated or full length. Reads that mapped between positions 130 and 134 were considered terminated and reads that mapped to position >=135 were considered antiterminated. Fraction terminator readthrough was calculated for each variant by dividing the number of antiterminated reads by the sum of terminated and antiterminated reads. Terminator efficiency is equal to 1 – fraction terminator readthrough.

### RNA structure prediction

RNA structure prediction was performed using the RNAstructure v6.1^36^ Fold command with default settings. For IH1 (Supplementary Fig. 2), WT sequences from previously identified ZTP riboswitches^26^ were obtained from the RefSeq database^57^ and aligned using INFERNAL v1.1.2^58^. The segment used for structure prediction contains sequence from the beginning of the multiple sequence alignment through the last unstructured nt within J1/2 (as determined by the RNAstructure Fold command^36^). Unbiased randomization allowed an equal probability for all nucleotides at each position. WT nucleotide distribution biased randomization was performed using the observed nucleotide frequency for each position as measured for the WT ZTP riboswitch multiple sequence alignment. All randomized data sets match the length distribution observed in the WT ZTP riboswitch set.

### L3n2 Nucleotide Frequency Analysis

P3 hairpin sequences from the multiple-sequence alignment described above that did not contain insertions were first binned by a match to the motif NNNNNGNCCNNNNGGGCNN and then binned by the number of predicted contiguous P3 base pairs. The frequency of each nucleotide at the position corresponding to nucleotide 90 of the *pfl* aptamer was then determined.

### Code Availability

Spats v1.0.1 can be accessed at https://github.com/LucksLab/spats/releases/. Scripts used in data processing are located at https://github.com/LucksLab/Cotrans_SHAPE-Seq_Tools/releases/ and https://github.com/LucksLab/LucksLab_Publications/tree/master/Strobel_ZTP_Riboswitch.

## Data Availability

Raw sequencing data that support the findings of this study have been deposited in the Sequencing Read Archive (SRA) (http://www.ncbi.nlm.nih.gov/sra) with the BioProject accession code <u>PRJNA510362</u>. Individual BioSample accession codes are available in Supplementary Table 1. SHAPE-Seq Reactivity Spectra generated in this work have been deposited in the RNA Mapping Database (RMDB) ^59^ (http://rmdb.stanford.edu/repository/) with the accession codes <u>ZTPRSW_BZCN_0001</u>, <u>ZTPRSW_BZCN_0002</u>, <u>ZTPRSW_BZCN_0003</u>, <u>ZTPRSW_BZCN_0004</u>, <u>ZTPRSW_BZCN_0005</u>, <u>ZTPRSW_BZCN_0006</u>, <u>ZTPRSW_BZCN_0007</u>, <u>ZTPRSW_BZCN_0008</u>, <u>ZTPRSW_BZCN_0009</u>, <u>ZTPRSW_BZCN_0010</u>, <u>ZTPRSW_BZCN_0011</u>, <u>ZTPRSW_BZCN_0012</u>, <u>ZTPRSW_BZCN_0013</u>, <u>ZTPRSW_BZCN_0014</u>, <u>ZTPRSW_BZCN_0015</u>, <u>ZTPRSW_BZCN_0016</u>. Sample details are available in Supplementary Table 2. Source data for all figures are available in the Northwestern University Arch Institutional Repository (https://doi.org/10.21985/N20F4H). All other data that support the findings of this paper are available from the corresponding authors upon request.

## Acknowledgments

We thank Rob Batey, Chad Torgerson, Scott Strobel, and Christopher Jones for thoughtful discussions; Jim Brink and Steve Hockema for review of combinatorial mutagenesis alignment software; Ronald Breaker and Keith Corbino for sharing a ZTP aptamer multiple sequence alignment; Kyle Watters for sharing a script to download RefSeq database entries. This work was supported by an Arnold O. Beckman Postdoctoral Fellowship (to E.J.S.), a New Innovator Award through the National Institute of General Medical Sciences of the National Institutes of Health (grant no. 1DP2GM110828 to J.B.L.), Searle Funds at The Chicago Community Trust (to J.B.L.) and by Grant Number T32GM008382 from the National Institute of General Medical Sciences. The content is solely the responsibility of the authors and does not necessarily represent the official views of the National Institutes of Health.

## Author Contributions

Eric J. Strobel, Conceptualization, Methodology, Investigation, Validation, Formal analysis, Data Curation, Software, Writing-original draft, Writing-review and editing, Supervision, Project Administration, Funding acquisition. Luyi Cheng, Investigation, Validation, Writing-review and editing. Katherine E. Berman, Investigation, Validation, Writing-review and editing. Paul D. Carlson, Investigation, Validation, Writing-review and editing. Julius B. Lucks, Methodology, Writing-original draft, Writing-reviewing and editing, Supervision, Resources, Project Administration, Funding acquisition.

## Supplementary Materials for

**Supplementary Figure 1.**
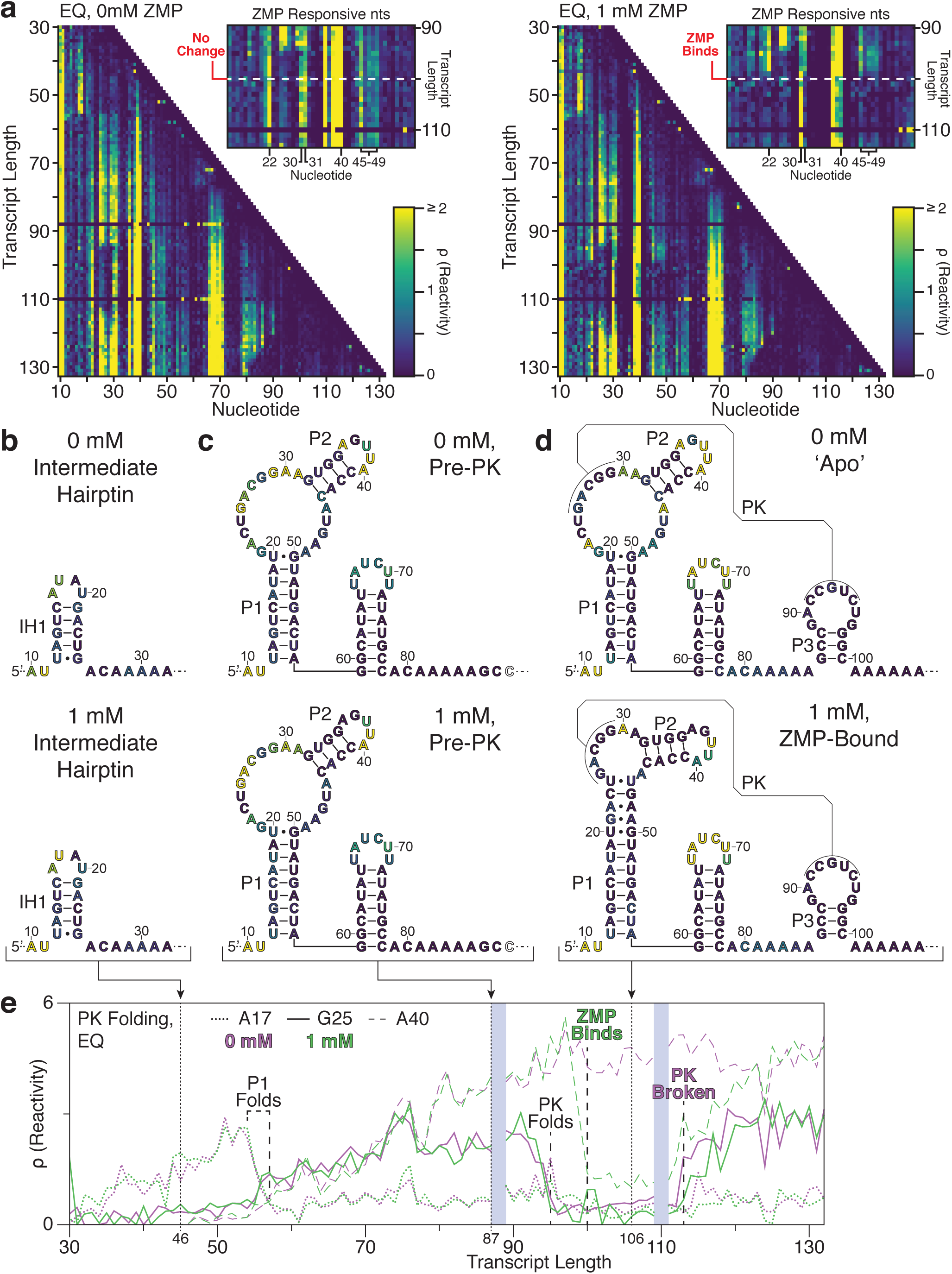
SHAPE probing of equilibrated *Cbe pfl* ZTP riboswitch intermediates. **(a)** Equilibrium refolded SHAPE-Seq reactivity matrix for the *Clostridium beijerinckii pfl* riboswitch with 0 mM and 1 mM ZMP. ZMP-responsive nts are highlighted in reactivity matrix cut-outs. The highly reactive unstructured leader (nts 1-9) is not shown and the absence of data for transcripts 88 and 110 is due to ambiguous alignment of 3’ ends. **(b)** Intermediate hairpin (IH1) secondary structure identified by manual analysis of reactivity values and refined with minimum free energy structure predictions of the leader region. Nucleotides are colored by the reactivity of transcript length 46 from (a). **(c)** Secondary structure of P1 and the linker region based, on the *pfl* RNA motif structure^26^. Nucleotides are colored by the reactivity of transcript length 87 from (a). **(d)** Apo and ZMP-bound secondary structure as determined by manual SHAPE-refinement of the consensus structure (Apo)^26^ or by the ZTP aptamer crystal structures^28–30^, respectively. Nucleotides are colored by the reactivity of transcript length 106 from (a). **(e)** ZTP riboswitch folding as depicted by reactivity trajectories for nucleotides A17 (within IH1 loop or P1 stem, dotted lines), G25 (within pseudoknot, solid lines), and A40 (within P2 loop, dashed lines) from (a). Trajectories are shown for both the absence (purple) and presence (green) of 1mM ZMP. Notable structural changes are identified by vertical dashed lines and the position of critical intermediate structures in (b-d) are shown by vertical dotted lines. Shaded boxes indicate the regions of poor alignment described in (a). Note that the annotated folding events are shifted to transcript lengths ~14 nts earlier relative to cotranscriptionally folded and probed transcripts in Figure 2 due to the absence of RNA polymerase. Results are for one experiment.

**Supplementary Figure 1-source data 1.** Source data for Supplementary Figure 1 are available in the Northwestern University Arch Institutional Repository (https://doi.org/10.21985/N20F4H).

**Supplementary Figure 2.**
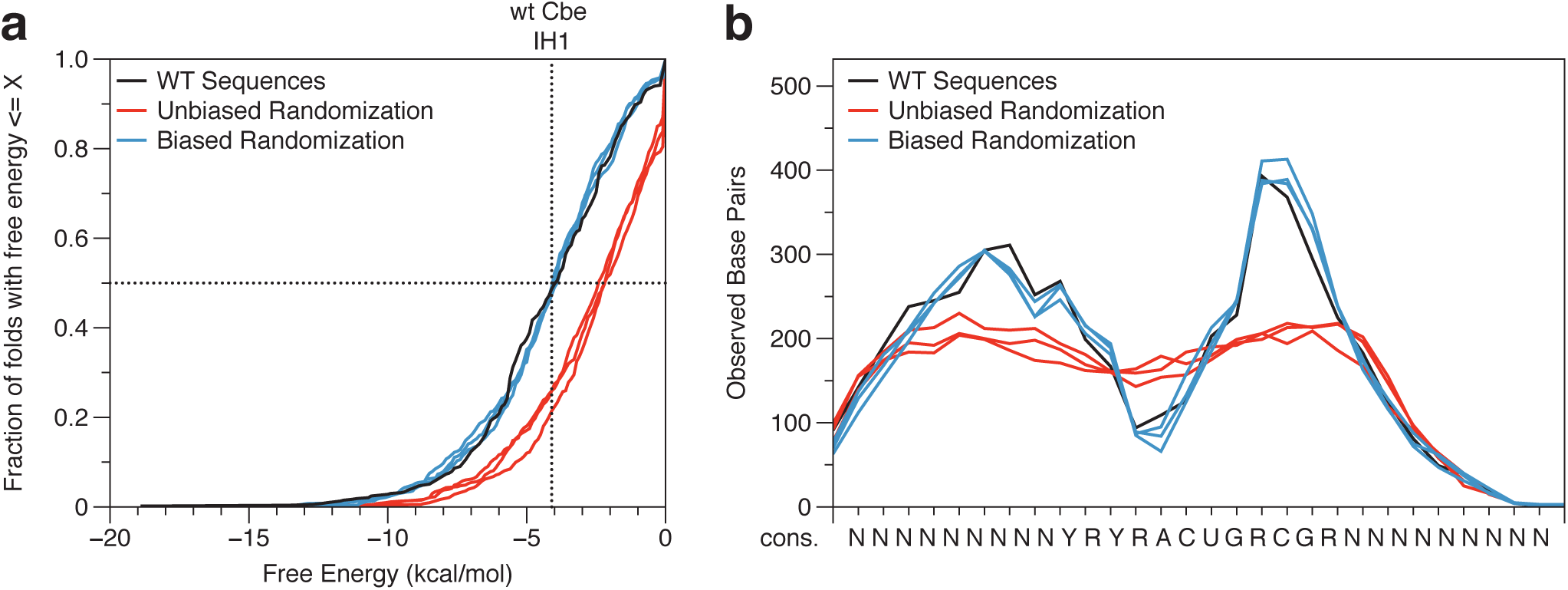
Analysis of ZTP riboswitch sequences for intermediate hairpin formation. **(a)** Cumulative distribution plot of the minimum free energy (ΔG) predicted for ZTP riboswitch sequences that correspond to the *pfl* riboswitch intermediate hairpin structure. Sequences comprise the first nucleotide in the multiple sequence alignment (9-10 nts upstream of J1/2) through the last unstructured nt within J1/2. The analysis of several sequence pools is presented: (black) 532 ZTP riboswitches from fully sequenced bacterial genomes^26^; (red) unbiased randomized sequences generated with an equal probability for observing each nucleotide at each position; (blue) sequences randomized by the natural nucleotide frequency at each position for the 532 sampled ZTP riboswitches^26^. All randomized data sets match the length distribution observed in the WT ZTP riboswitch set. **(b)** The number of base pairs predicted at each position as observed in the structures predicted in (a).

**Supplementary Figure 2-source data 2.** Source data for Supplementary Figure 2 are available in the Northwestern University Arch Institutional Repository (https://doi.org/10.21985/N20F4H).

**Supplementary Figure 3.**
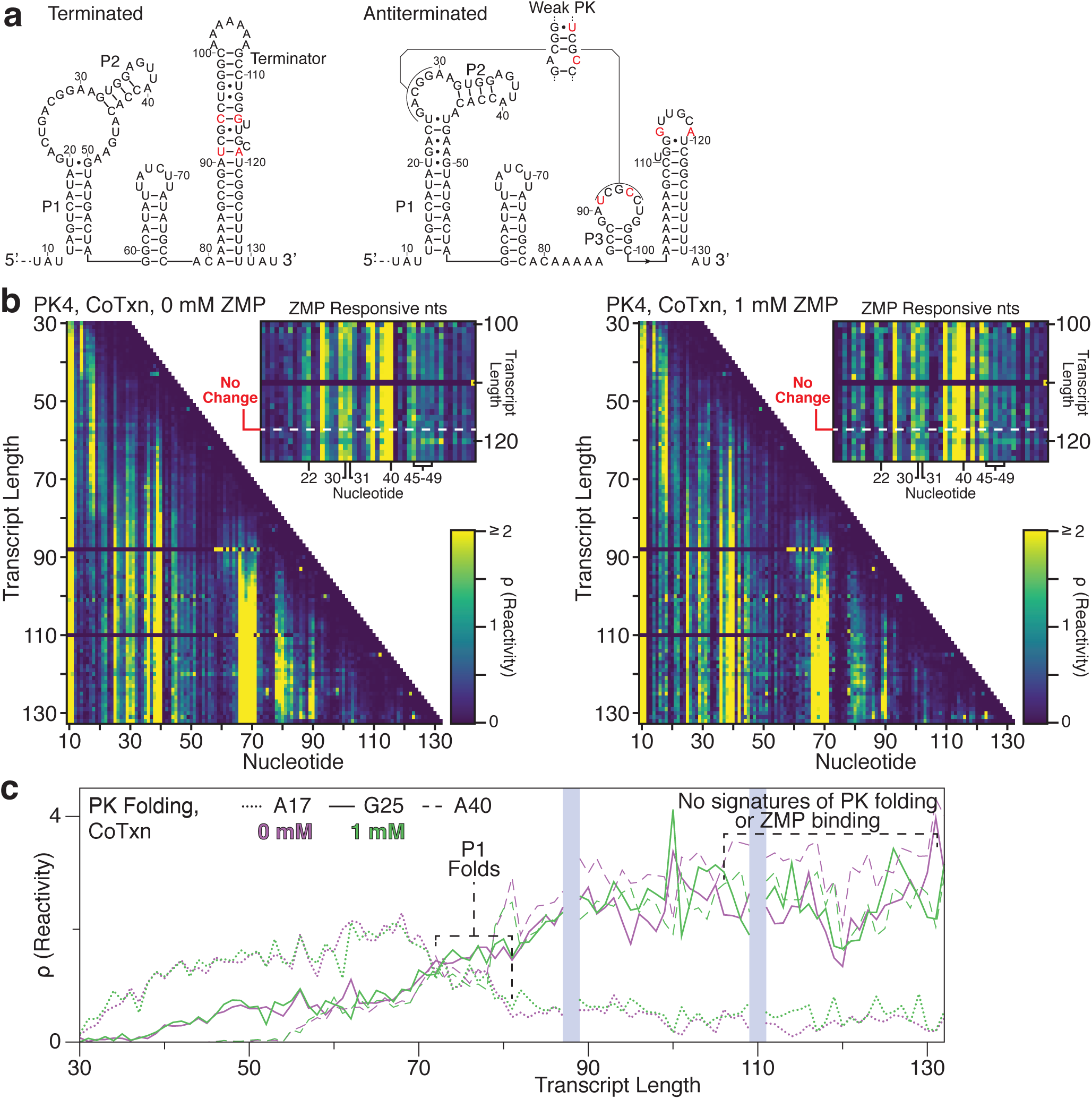
Cotranscriptional SHAPE probing of disrupted pseudoknot mutant. **(a)** *Clostridium beijerinckii pfl* riboswitch terminated and antiterminated secondary structures shown with PK4 (C91U,U94C,A114G,G119A) mutations. The base pair configuration of the mutated pseudoknot is shown. **(b)** Cotranscriptional SHAPE-Seq reactivity matrix for the *Clostridium beijerinckii* riboswitch PK4 mutant with 0 mM and 1 mM ZMP. ZMP-responsive nts are highlighted in reactivity matrix cut-outs. The highly reactive unstructured leader (nts 1-9) is not shown and the absence of data for transcripts 88 and 110 is due to ambiguous alignment of 3’ ends. **(c)** ZTP riboswitch folding as depicted by reactivity trajectories for nucleotides A17 (within IH1 loop or P1 stem, dotted lines), G25 (within pseudoknot, solid lines), and A40 (within P2 loop, dashed lines) from (a). Trajectories are shown for both the absence (purple) and presence (green) of 1mM ZMP. Notable structural changes are identified by vertical dashed lines. Shaded boxes indicate the regions of poor alignment described in (a). Results are for one experiment.

**Supplementary Figure 3-Source data.** Source data for Supplementary Figure 3 are available in the Northwestern University Arch Institutional Repository (https://doi.org/10.21985/N20F4H).

**Supplementary Figure 4.**
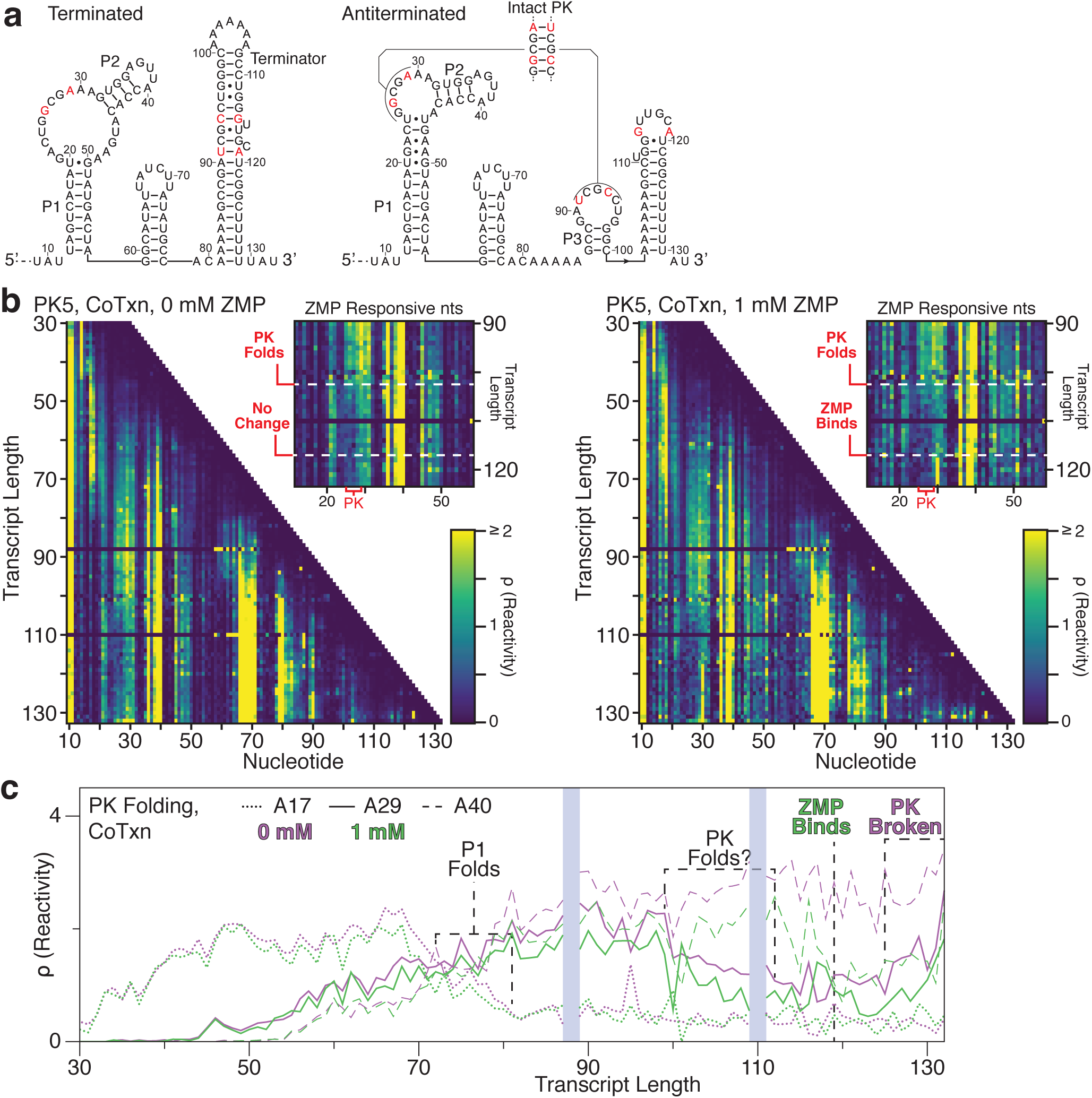
Cotranscriptional SHAPE probing of restored pseudoknot mutant. **(a)** *Clostridium beijerinckii* riboswitch terminated and antiterminated secondary structures shown with PK5 (A26G,G29A,C91U,U94C,A114G,G119A) mutations. The base pair configuration of the mutated pseudoknot is shown. **(b)** Cotranscriptional SHAPE-Seq reactivity matrix for the *Clostridium beijerinckii* riboswitch PK5 mutant with 0 mM and 1 mM ZMP. ZMP-responsive nts are highlighted in reactivity matrix cut-outs. The highly reactive unstructured leader (nts 1-9) is not shown and the absence of data for transcripts 88 and 110 is due to ambiguous alignment of 3’ ends. **(c)** ZTP riboswitch folding as depicted by reactivity trajectories for nucleotides A17 (within IH1 loop or P1 stem, dotted lines), A29 (within pseudoknot, solid lines), and A40 (within P2 loop, dashed lines) from (a). For PK5, pseudoknot nucleotides from 25-29 were more evenly reactive than the WT sequence and A29 is shown as a representative for clarity. Trajectories are shown for both the absence (purple) and presence (green) of 1mM ZMP. Notable structural changes are identified by vertical dashed lines. Shaded boxes indicate the regions of poor alignment described in (a). Results are for one experiment.

**Supplementary Figure 4-Source data.** Source data for Supplementary Figure 4 are available in the Northwestern University Arch Institutional Repository (https://doi.org/10.21985/N20F4H).

**Supplementary Figure 5.**
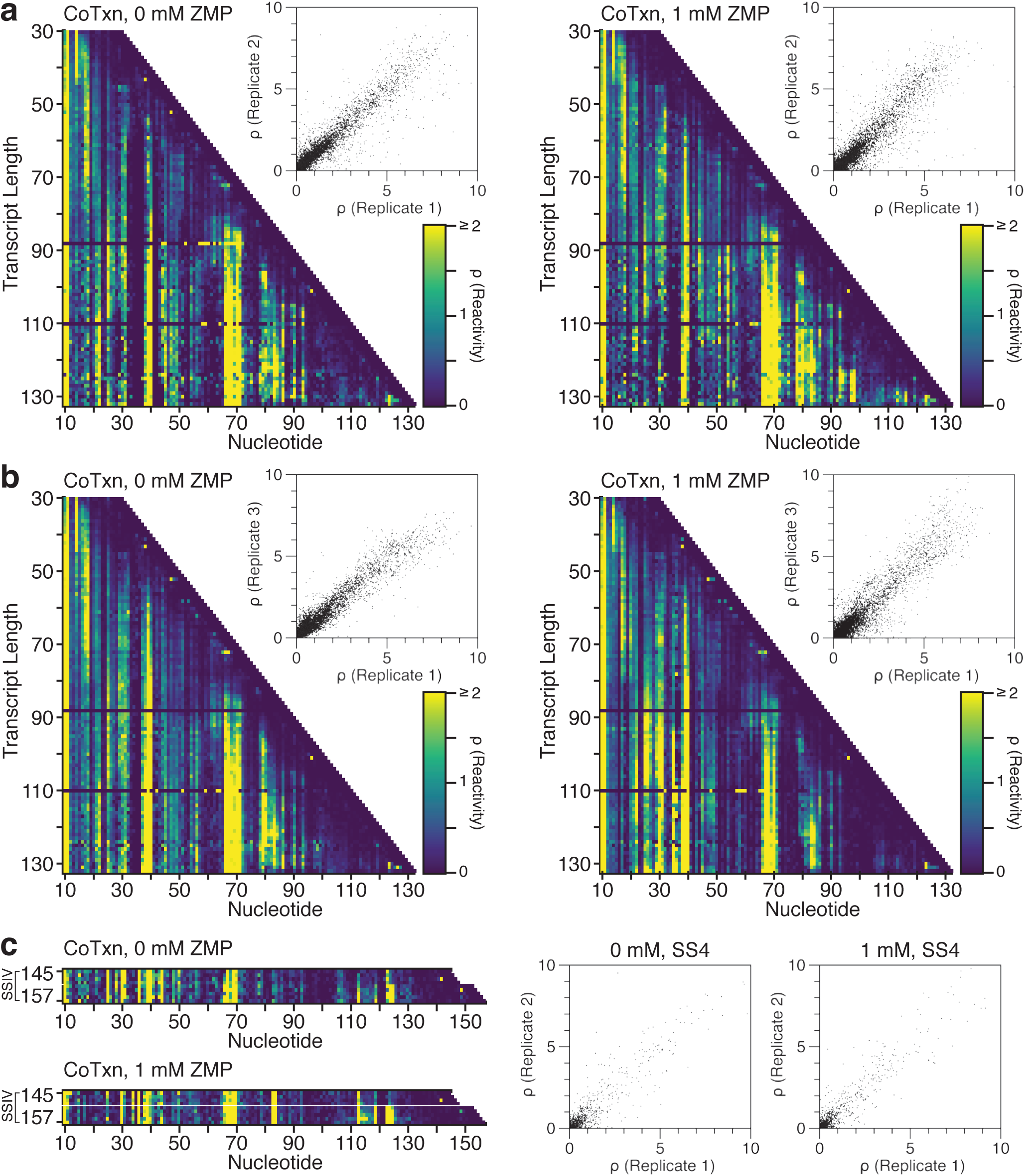
Cotranscriptional SHAPE probing of the *Cbe pfl* ZTP riboswitch Replicate Data. **(a)** Cotranscriptional SHAPE-Seq reactivity matrix for the *Clostridium beijerinckii pfl* ZTP riboswitch with 0 mM and 1 mM ZMP; Replicate 2. The highly reactive unstructured leader (nts 1-9) is not shown and the absence of data for transcripts 88 and 110 is due to ambiguous alignment of 3’ ends. Reactivity values from replicate 2 transcript lengths 30 to 132 are plotted against the corresponding values from replicate 1 for each condition. **(b)** As in (a), but for replicate 3. **(c)** Cotranscriptional SHAPE-Seq reactivity matrix for the *Clostridium beijerinckii pfl* ZTP riboswitch with 0 mM and 1 mM ZMP targeting transcripts 145-148 and 153-157 using terminal biotin roadblocks and Superscript IV reverse transcriptase (RT); Replicate 2. Reactivity values from replicate 2 transcript lengths 30 to 132 are plotted against the corresponding values from replicate 1 for each condition.

**Supplementary Figure 5-Source data.** Source data for Supplementary Figure 5 are available in the Northwestern University Arch Institutional Repository (https://doi.org/10.21985/N20F4H).

**Supplementary Figure 6.**
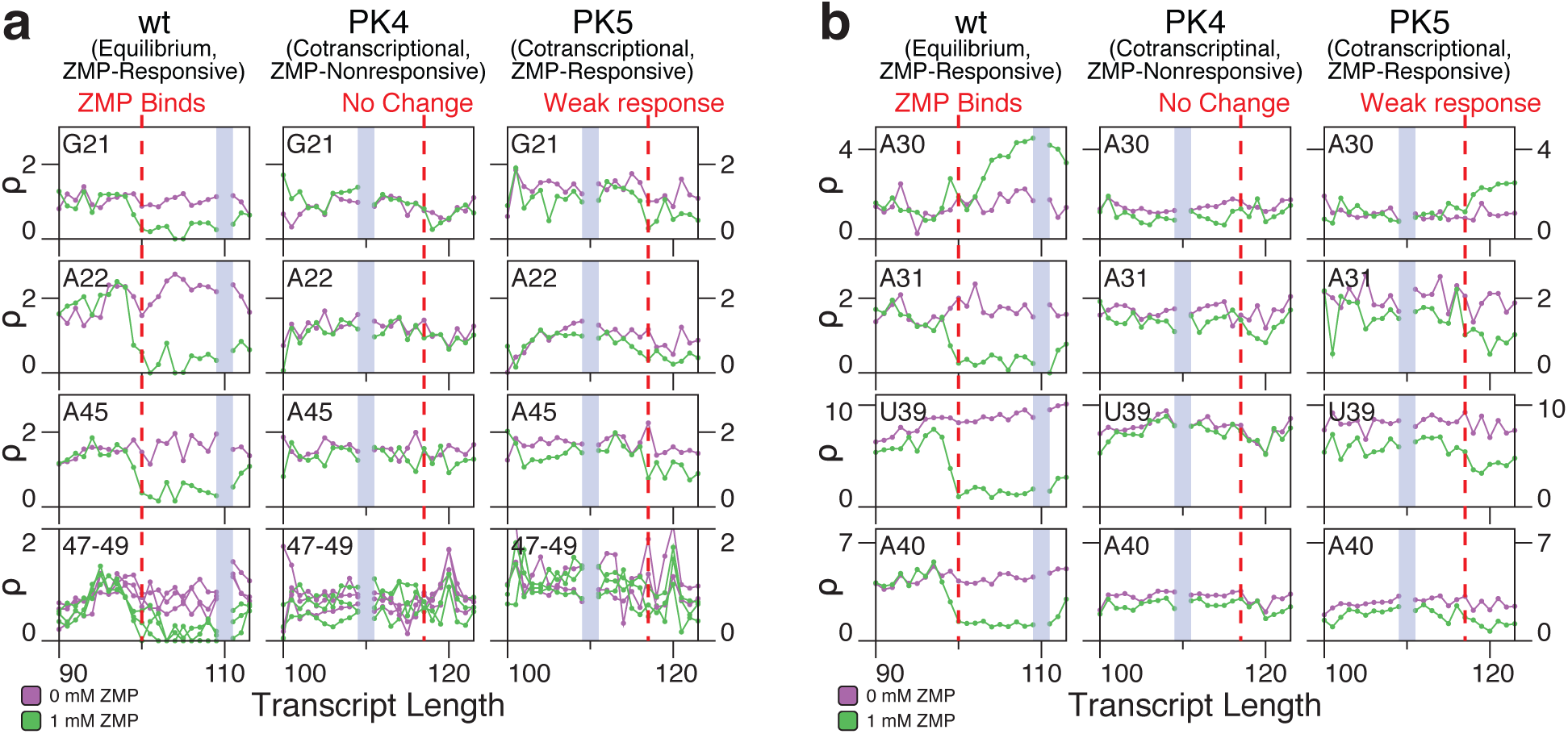
SHAPE reactivity of ZMP-responsive nucleotides in equilibrated WT and cotranscriptionally folded pseudoknot mutants. ZMP-responsive nucleotides within the *Cbe pfl* ZTP riboswitch, including **(a)** non-canonical P1 base pairs and **(b)** the nucleotides that interact with the pseudoknot as defined in Figure 3. SHAPE-Seq reactivity traces across transcript lengths are shown in the absence (purple) and presence (green) of 1mM ZMP. WT equilibrium refolded data are from Supplementary Figure 1 and PK4/PK5 cotranscriptionally folded data are from Supplementary Figures 3 and 4. The vertical red dashed lines indicate the transcript length at which ZMP binding is inferred to occur based on bifurcation of the WT trajectories. Results are for one experiment.

**Supplementary Figure 6-source data.** Source data for Supplementary Figure 6 are available in the Northwestern University Arch Institutional Repository (https://doi.org/10.21985/N20F4H).

**Supplementary Figure 7.**
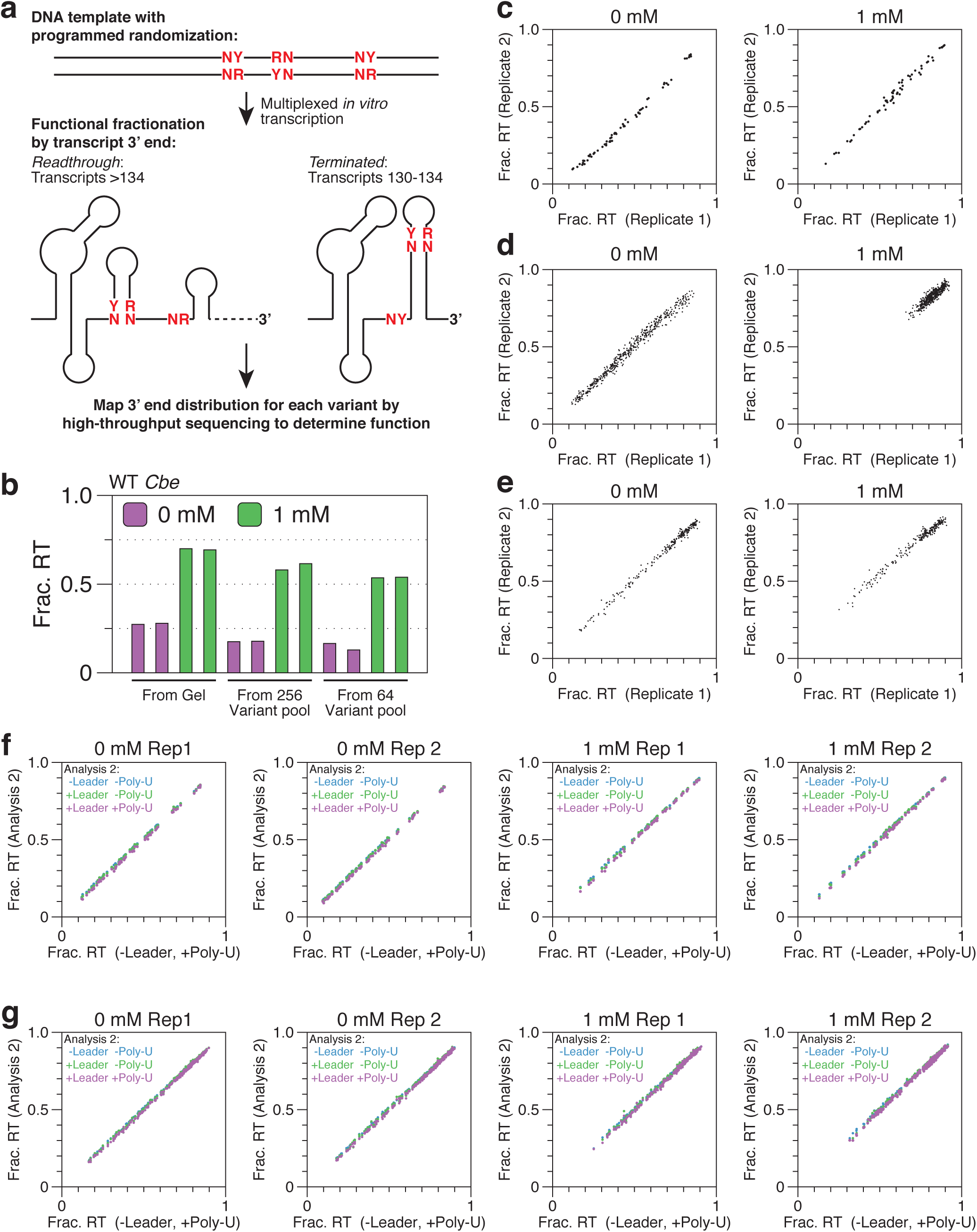
Targeted *in vitro* transcription and data analysis controls. **(a)** Overview of combinatorial mutagenesis experiment. A linear DNA template library is constructed using a long synthetic oligonucleotide with randomization at specified positions. Multiplexed *in vitro* transcription is performed using this DNA template library to produce a distribution of 3’ ends for each variant within the library. High-throughput sequencing is then used to measure this distribution for every variant to determine its function. **(b)** Fraction readthrough for the WT *Cbe* ZTP riboswitch with 0 mM and 1 mM as measured by targeted *in vitro* transcription and gel electrophoresis (data from Fig. 1c) or by high-throughput sequencing as part of a combinatorial mutagenesis library with 256 (data from experiment in Supplementary Figure 11) or 64 (data from experiment in Figure 4) variants. **(c)** Plot of fraction readthrough values for all aptamer/terminator overlap combinatorial mutagenesis variants from replicate 1 against replicate 2 for 0 mM and 1 mM ZMP conditions (data from Figure 4). **(d)** Plot of fraction readthrough values for all P3 stem combinatorial mutagenesis variants from replicate 1 against replicate 2 for 0 mM and 1 mM ZMP conditions (data from Figure 5). **(e)** Plot of fraction readthrough values for all pseudoknot contact combinatorial mutagenesis variants from replicate 1 against replicate 2 for 0 mM and 1 mM ZMP conditions (data from Supplementary Figure 11). **(f)** Analysis controls in which sequencing alignment omitted the ZTP riboswitch 5’ leader, poly-U tract, or both. Each analysis control plotted against alignment without the 5’ leader and with the poly-U tract. Data are from Figure 4. **(g)** As in (f) but data are from Supplementary Figure 11.

**Supplementary Figure 7-source data.** Source data for Supplementary Figure 7 are available in the Northwestern University Arch Institutional Repository (https://doi.org/10.21985/N20F4H).

**Supplementary Figure 8.**
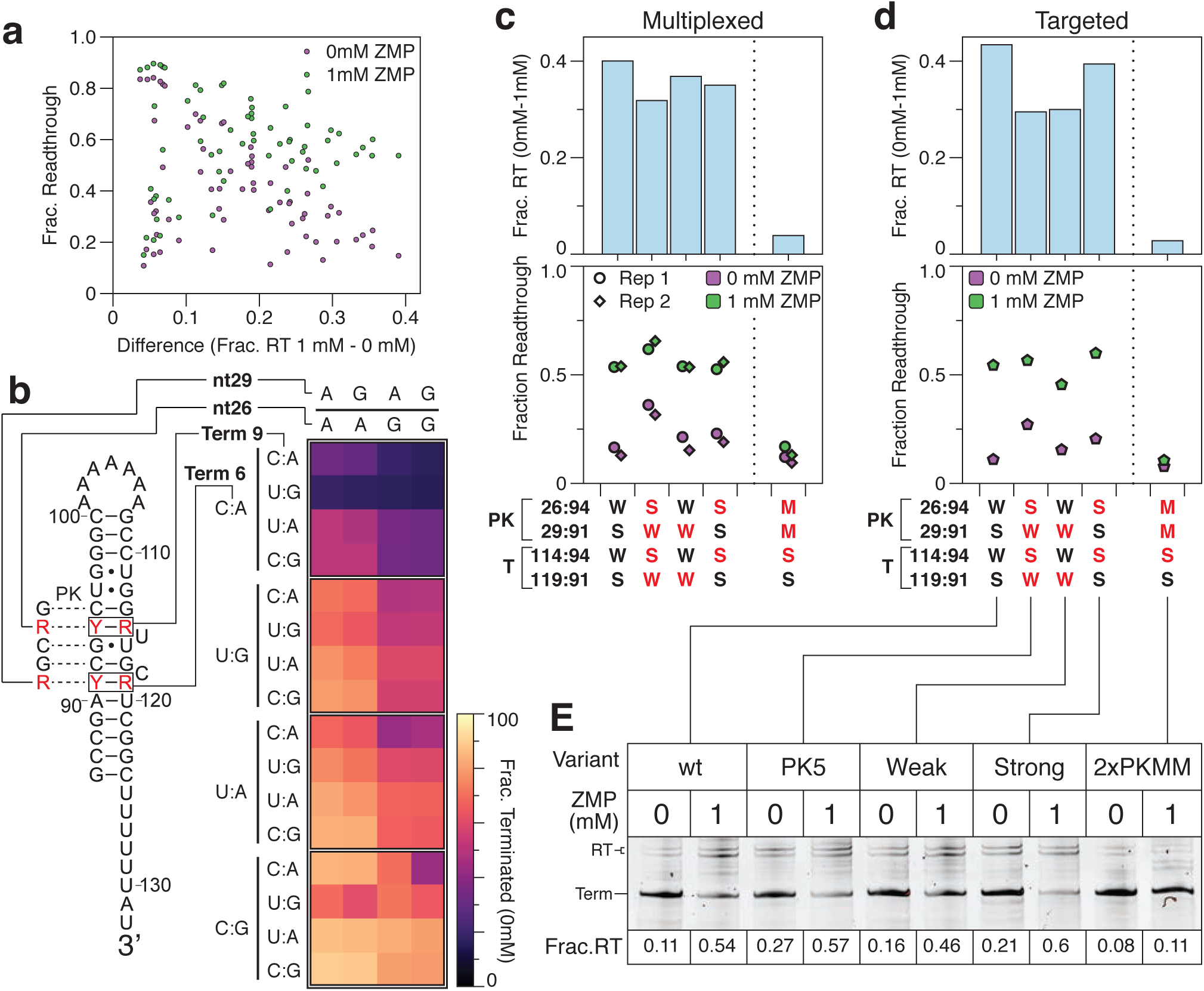
Complete aptamer/terminator overlap combinatorial mutagenesis data. **(a)** Plot of fraction readthrough of aptamer/terminator overlap combinatorial mutagenesis mutants from Figure 4a as measured in the absence (purple) and presence (green) of 1 mM ZMP ordered by the difference in fraction readthrough observed in 1 mM and 0 mM ZMP conditions. **(b)** Terminator efficiency (1 – fraction readthrough) of aptamer/terminator overlap combinatorial mutagenesis mutants from Figure 4a in the absence of ZMP. Mutants are grouped vertically by the inferred stability of terminator base pairs 91:119 (Term 6) and 94:114 (Term 9) and horizontally by the inferred stability of pseudoknot base pairs 26:94 and 29:91. **(c)** Fraction readthrough for mutants with complete pseudoknot and terminator base pair matches and for a mutant with a complete terminator match and pseudoknot mismatch. The average difference in fraction readthrough (1 mM – 0 mM) is shown above each variant. Variant pairing patterns in the pseudoknot (PK) and terminator (T) are annotated as weak (W, A-U), strong (S, G-C), and mismatch (M, A-C). Red annotations indicate deviations from the wild-type pairing pattern. **(d)** Fraction readthrough for mutants from (c) as measured by targeted *in vitro* transcription and gel electorophoresis. **(e)** *in vitro* transcription gel for measurements in (d). n=2 independent biological replicates are annotated as ‘Rep 1’ and ‘Rep 2’ in panel c. Panels a and b are the average of n = 2 independent biological replicates. Individual replicate values are compared in Supplementary Figure 7c. Panels d and e are from one experiment.

**Supplementary Figure 8-source data.** Source data for Supplementary Figure 8 are available in the Northwestern University Arch Institutional Repository (https://doi.org/10.21985/N20F4H).

**Supplementary Figure 9.**
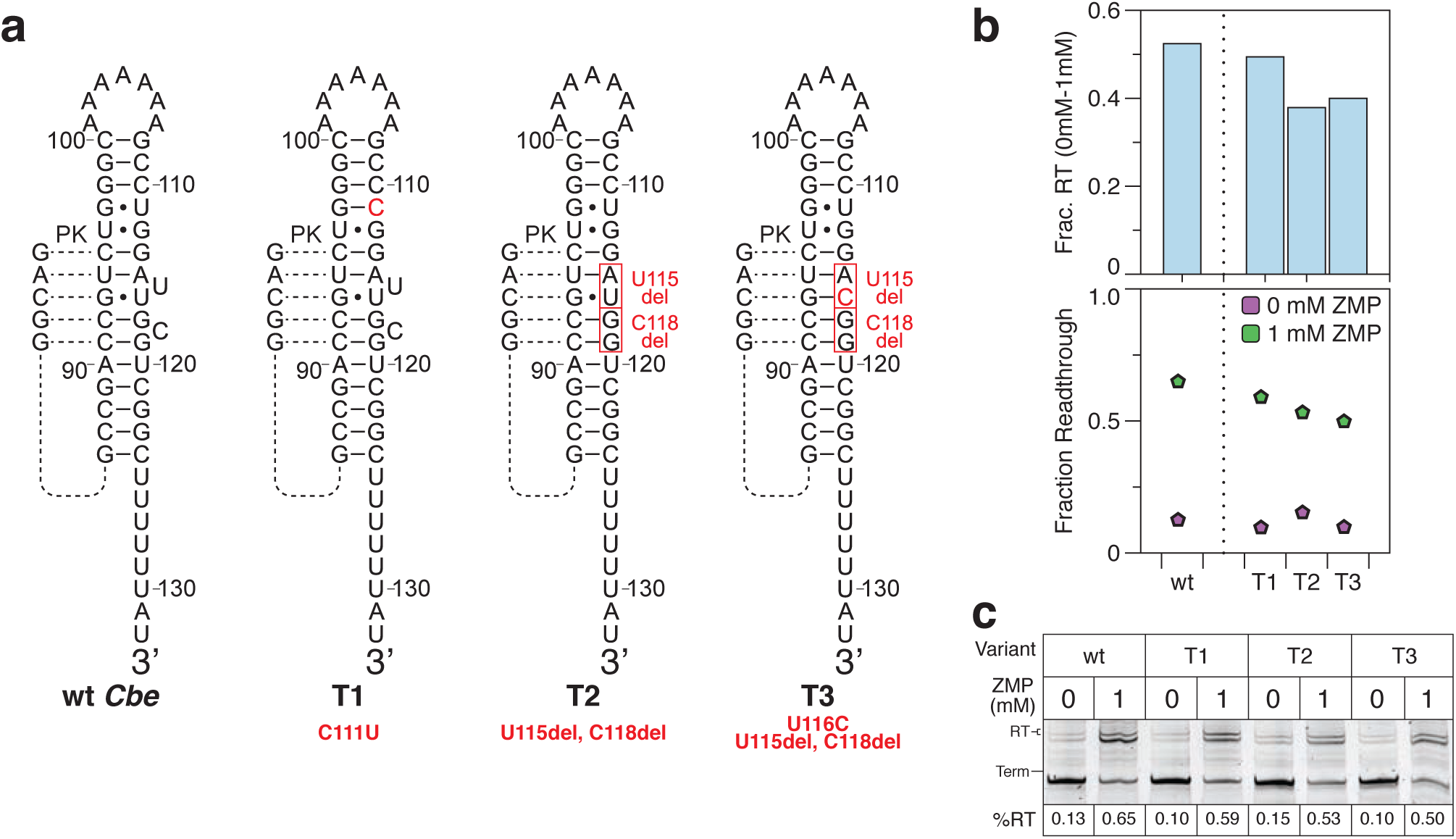
*in vitro* transcription of *Cbe* ZTP riboswitch terminator variants. **(a)** Expected secondary structure of intrinsic terminator hairpin for mutants analyzed in (b). The position of pseudoknot base pairs that compete with the formation of the 3’ terminator stem is shown. **(b)** Fraction readthrough for terminator mutants shown in (A) as measured in the absence (purple) and presence (green) of 1 mM ZMP. The difference in fraction readthrough (1 mM – 0 mM) is shown above each variant. **(c)** *in vitro* transcription gel for measurements in (b). Results are from one experiment.

**Supplementary Figure 9-source data.** Source data for Supplementary Figure 9 are available in the Northwestern University Arch Institutional Repository (https://doi.org/10.21985/N20F4H).

**Supplementary Figure 10.**
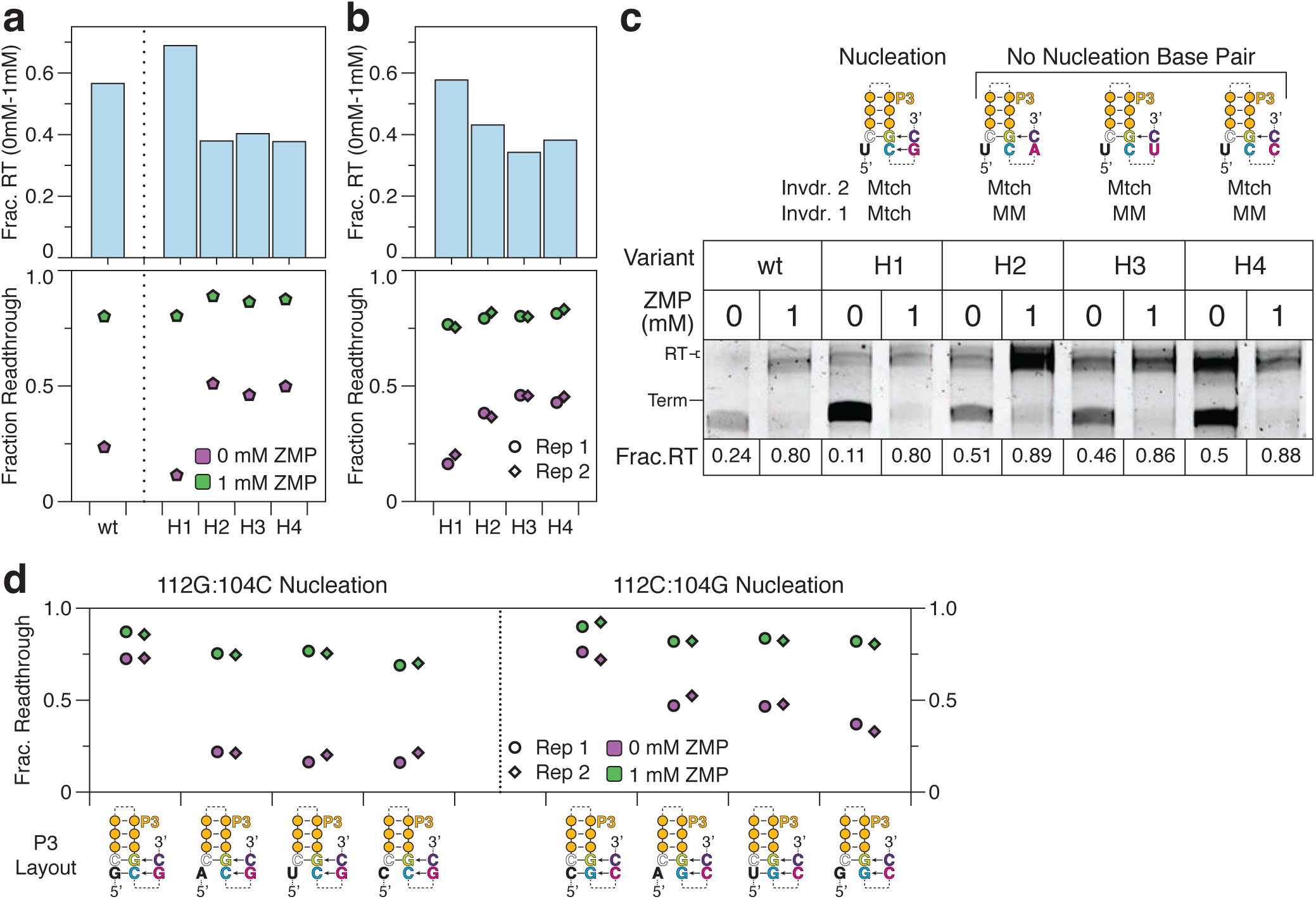
*in vitro* transcription of *Cbe pfl* ZTP riboswitch terminator nucleation variants. **(a)** Targeted *in vitro* transcription of WT *Cbe pfl* riboswitch and terminator nucleation variants. Fraction readthrough for variants as measured in the absence (purple) and presence (green) of 1 mM ZMP. The difference in fraction readthrough (1 mM – 0 mM) is shown above each variant. The configuration for terminator strand displacement of P3 for each variant is shown in (c). **(b)** Fraction readthrough for variants in (a) as measured by combinatorial mutagenesis. **(c)** *In* vitro transcription gel for the measurements in (a). The configuration for terminator strand displacement of P3 for each variant is shown as in Figure 5. **(d)** Fraction readthrough for select variants with terminator hairpin nucleation by 112G:104C or 112:C104G base pairs. Results in (a) are from one experiment. Results in (b) and (d) are from n=2 independent biological replicates; the difference in fraction readthrough (0mM-1mM) is the average of these replicates.

**Supplementary Figure 10 source data.** Source data for Supplementary Figure 10 are available in the Northwestern University Arch Institutional Repository (https://doi.org/10.21985/N20F4H).

**Supplementary Figure 11.**
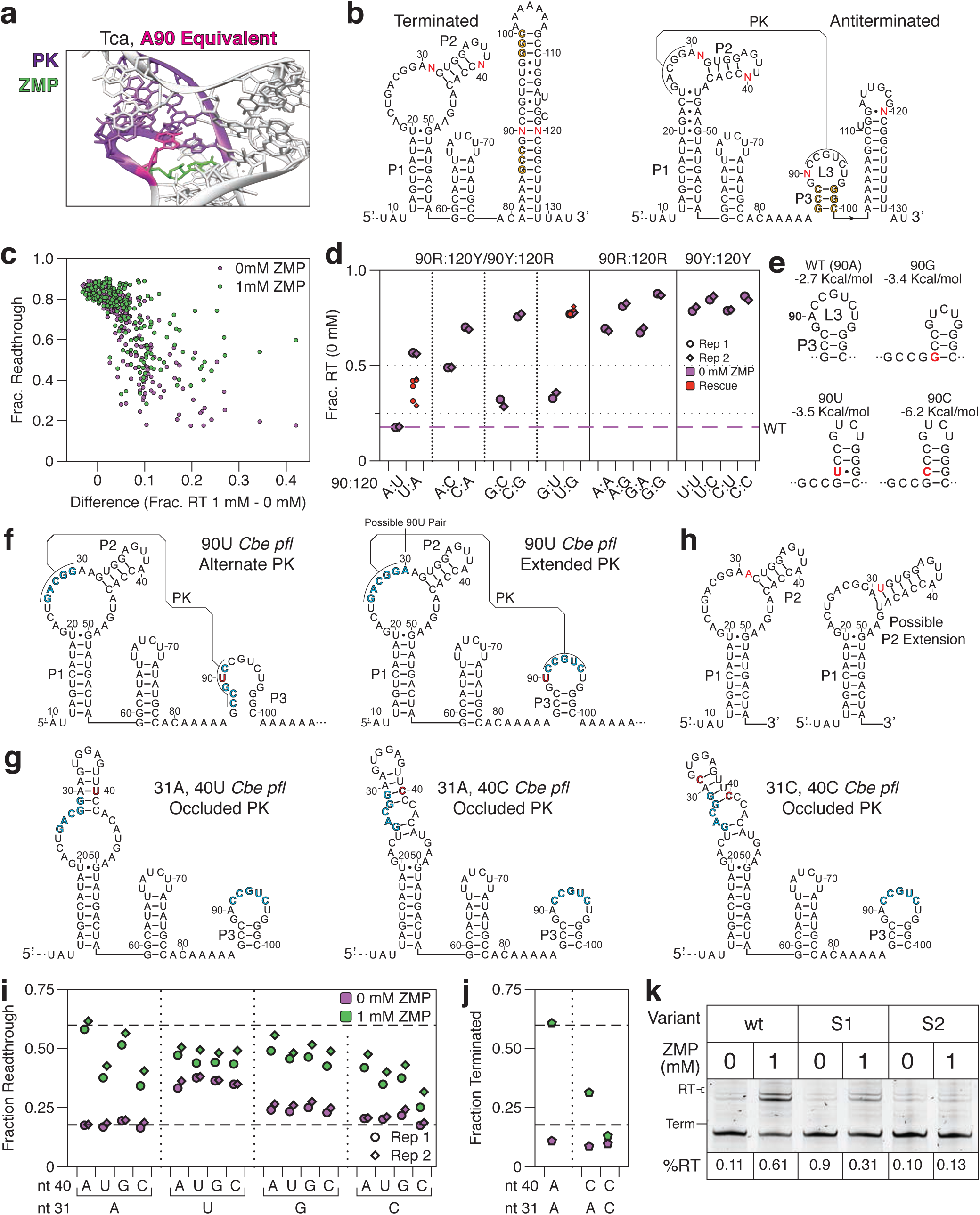
Combinatorial mutagenesis of pseudoknot contact variants. **(a)** The crystal structure of the *Thermosinus carboxydivorans* ZMP riboswitch (PDB: 4ZNP)^30^ is highlighted to show the interaction between the nucleotide equivalent to *Cbe pfl* A90 (magenta) interacting with the Hoogsteen edge of a pseudoknot nucleotide (purple). **(b)** *Cbe* ZTP riboswitch terminated and antiterminated secondary structures depicting the randomization scheme used for combinatorial mutagenesis of nucleotides that contact the ZTP aptamer psuedoknot. P3 stem nucleotides are gold and randomized nucleotides are red. **(c)** Plot of fraction readthrough of combinatorial mutagenesis mutants from (b) as measured in the absence (purple) and presence (green) of 1 mM ZMP ordered by the difference in fraction readthrough observed in 1 mM and 0 mM ZMP conditions. **(d)** Fraction readthrough in the absence of ZMP for variants from (b) grouped by the identity of nucleotides 90 and 120; Large purple points contain the WT nucleotides 31A and 40A; Small red points indicate ‘rescue’ mutants (31A, 40U; 31A, 40C; 31C, 40C) that are predicted to partially occlude the pseudoknot. **(e)** Structure predictions for each N90 variant of the P3 region (nts 86-100) with an associated ΔG predicted by the RNAstructure Fold command. **(f)** Possible misfolded pseudoknot configurations due to a 90U mutation. Pseudoknot nucleotides are shown in blue and mutations are shown in red. **(g)** Minimum free energy structure predictions for the 31A,40U; 31A,40C; and 31C40C variants as predicted by the RNAstructure Fold command. The predicted structures are supported by the observation that the 31A:40Y defect is rescued by 31A to 31G mutations that convert an A:U pair in misfolded structures to G:U (compare panel (i) 31A mutants to 31G mutants), and the 31C,40C mutation is rescued by all 31D,40C combinations (D = A, G, U) which disrupt a G:C pair in the misfolded structure (panel (i), 40C mutants). **(h)** Secondary structure of the WT *Cbe* ZTP aptamer and an extended P2 variation that could occur for A31U mutants. The 31U mutation is shown in red. **(i)** Fraction readthrough for nucleotide 31 and 40 mutants with the WT terminator (90A, 120U). Dashed lines indicate fraction readthrough observed for the WT *Cbe* ZTP riboswitch in each condition. **(j)** Targeted *in vitro* transcription analysis of select mutants from (i). **(k)** *In vitro* transcription gel for measurements in (j). n=2 independent biological replicates are annotated as ‘Rep 1’ and ‘Rep 2’ in panel d and i. Panel c (Frac. Readthrough 1mM-0mM) is the average of n=2 independent biological replicates. Individual replicate values are compared in Supplementary Figure 7. Panels j and k are from one experiment.

**Supplementary Figure 11-source data.** Source data for Supplementary Figure 11 are available in the Northwestern University Arch Institutional Repository (https://doi.org/10.21985/N20F4H).

**Supplementary Table 1.** Sequencing Read Archive (SRA) deposition table. All primary sequencing data generated in this work are freely available from the Sequencing Read Archive (http://www.ncbi.nlm.nih.gov/sra), accessible via the BioProject accession number <u>PRJNA510362</u> or using the individual accession numbers below.

| **SRA Accession** | **RNA** | **Experiment** | **Figure(s)** |
| --- | --- | --- | --- |
| SAMN10607306 | ZTP riboswitch, WT (rep.1) | 0 mM ZMP, cotranscriptional | Figs. 2, 3 |
| SAMN10607307 | ZTP riboswitch, WT (rep. 2) | 0 mM ZMP, cotranscriptional | Supp. Fig. 5 |
| SAMN10607308 | ZTP riboswitch, WT (rep. 3) | 0 mM ZMP, cotranscriptional | Supp. Fig. 5 |
| SAMN10607309 | ZTP riboswitch, WT (rep. 1) | 1 mM ZMP, cotranscriptional | Figs. 2, 3 |
| SAMN10607310 | ZTP riboswitch, WT (rep. 2) | 1 mM ZMP, cotranscriptional | Supp. Fig. 5 |
| SAMN10607311 | ZTP riboswitch, WT (rep. 3) | 1 mM ZMP, cotranscriptional | Supp. Fig. 5 |
| SAMN10607312 | ZTP riboswitch, WT (SSIV rep. 1) | 0 mM ZMP, cotrans, SSIV RT | Supp. Fig. 5 |
| SAMN10607313 | ZTP riboswitch, WT (SSIV rep. 2) | 0 mM ZMP, cotrans, SSIV RT | Fig. 2 |
| SAMN10607314 | ZTP riboswitch, WT (SSIV rep. 1) | 1 mM ZMP, cotrans, SSIV RT | Supp. Fig. 5 |
| SAMN10607315 | ZTP riboswitch, WT (SSIV rep. 2) | 1 mM ZMP, cotrans, SSIV RT | Fig. 2 |
| SAMN10607316 | ZTP riboswitch, WT | 0 mM ZMP, equilibrium | Supp. Fig. 1<br>Supp. Fig. 6 |
| SAMN10607317 | ZTP riboswitch, WT | 1 mM ZMP, equilibrium | Supp. Fig. 1<br>Supp. Fig. 6 |
| SAMN10607318 | ZTP riboswitch, PK4 Mutant | 0 mM ZMP, cotranscriptional | Supp. Fig. 3 |
| SAMN10607319 | ZTP riboswitch, PK4 Mutant | 1 mM ZMP, cotranscriptional | Supp. Fig. 3 |
| SAMN10607320 | ZTP riboswitch, PK5 Mutant | 0 mM ZMP, cotranscriptional | Supp. Fig. 4 |
| SAMN10607321 | ZTP riboswitch, PK5 Mutant | 1 mM ZMP, cotranscriptional | Supp. Fig. 4 |
| SAMN10607322 | ZTP RndPK (rep. 1) | 0 mM ZMP | Fig. 4<br>Supp. Fig. 7<br>Supp. Fig. 8 |
| SAMN10607323 | ZTP RndPK (rep. 2) | 0 mM ZMP | Fig. 4<br>Supp. Fig. 7<br>Supp. Fig. 8 |
| SAMN10607324 | ZTP RndPK (rep. 1) | 1 mM ZMP | Fig. 4<br>Supp. Fig. 7<br>Supp. Fig. 8 |
| SAMN10607325 | ZTP RndPK (rep. 2) | 1 mM ZMP | Fig. 4<br>Supp. Fig. 7<br>Supp. Fig. 8 |
| SAMN10607326 | ZTP RndP3v2 (rep. 1) | 0 mM ZMP | Fig. 5<br>Supp. Fig. 7<br>Supp. Fig. 10 |
| SAMN10607327 | ZTP RndP3v2 (rep. 2) | 0 mM ZMP | Fig. 5<br>Supp. Fig. 7<br>Supp. Fig. 10 |
| SAMN10607328 | ZTP RndP3v2 (rep. 1) | 1 mM ZMP | Fig. 5<br>Supp. Fig. 7<br>Supp. Fig. 10 |
| SAMN10607329 | ZTP RndP3v2 (rep. 2) | 1 mM ZMP | Fig. 5<br>Supp. Fig. 7<br>Supp. Fig. 10 |
| SAMN10607330 | ZTP RndPKSTB (rep. 1) | 0 mM ZMP | Fig. 6<br>Supp. Fig. 7<br>Supp. Fig. 11 |
| SAMN10607331 | ZTP RndPKSTB (rep. 2) | 0 mM ZMP | Fig. 6<br>Supp. Fig. 7<br>Supp. Fig. 11 |
| SAMN10607332 | ZTP RndPKSTB (rep. 1) | 1 mM ZMP | Fig. 6<br>Supp. Fig. 7<br>Supp. Fig. 11 |
| SAMN10607333 | ZTP RndPKSTB (rep. 2) | 1 mM ZMP | Fig. 6<br>Supp. Fig. 7<br>Supp. Fig. 11 |

**Supplementary Table 2.** RMDB data deposition table. SHAPE-Seq reactivity spectra generated in this work are freely available from the RNA Mapping Database (RMDB) (http://rmdb.stanford.edu/repository/)^59^, accessible using the RMDB ID numbers indicated in the table below.

| **RMDB Accession** | **RNA** | **Experiment** | **Figure(s)** |
| --- | --- | --- | --- |
| ZTPRSW_BZCN_0001 | ZTP riboswitch, WT (rep.1) | 0 mM ZMP, cotranscriptional | Figs. 2, 3 |
| ZTPRSW_BZCN_0002 | ZTP riboswitch, WT (rep. 2) | 0 mM ZMP, cotranscriptional | Supp. Fig. 5 |
| ZTPRSW_BZCN_0003 | ZTP riboswitch, WT (rep. 3) | 0 mM ZMP, cotranscriptional | Supp. Fig. 5 |
| ZTPRSW_BZCN_0004 | ZTP riboswitch, WT (rep. 1) | 1 mM ZMP, cotranscriptional | Figs. 2, 3 |
| ZTPRSW_BZCN_0005 | ZTP riboswitch, WT (rep. 2) | 1 mM ZMP, cotranscriptional | Supp. Fig. 5 |
| ZTPRSW_BZCN_0006 | ZTP riboswitch, WT (rep. 3) | 1 mM ZMP, cotranscriptional | Supp. Fig. 5 |
| ZTPRSW_BZCN_0011 | ZTP riboswitch, WT | 0 mM ZMP, equilibrium | Supp. Fig.1<br>Supp. Fig. 6 |
| ZTPRSW_BZCN_0012 | ZTP riboswitch, WT | 1 mM ZMP, equilibrium | Supp. Fig.1<br>Supp. Fig. 6 |
| ZTPRSW_BZCN_0007 | ZTP riboswitch, WT (SSIV rep. 1) | 0 mM ZMP, cotranscriptional,<br>SSIV RT | Supp. Fig. 5 |
| ZTPRSW_BZCN_0008 | ZTP riboswitch, WT (SSIV rep. 2) | 0 mM ZMP, cotranscriptional,<br>SSIV RT | Fig. 2 |
| ZTPRSW_BZCN_0009 | ZTP riboswitch, WT (SSIV rep. 1) | 1 mM ZMP, cotranscriptional,<br>SSIV RT | Supp. Fig. 5 |
| ZTPRSW_BZCN_0010 | ZTP riboswitch, WT (SSIV rep. 2) | 1 mM ZMP, cotranscriptional,<br>SSIV RT | Fig. 2 |
| ZTPRSW_BZCN_0013 | ZTP riboswitch, PK4 Mutant | 0 mM ZMP, cotranscriptional | Supp. Fig. 3 |
| ZTPRSW_BZCN_0014 | ZTP riboswitch, PK4 Mutant | 1 mM ZMP, cotranscriptional | Supp. Fig. 3 |
| ZTPRSW_BZCN_0015 | ZTP riboswitch, PK5 Mutant | 0 mM ZMP, cotranscriptional | Supp. Fig. 4 |
| ZTPRSW_BZCN_0016 | ZTP riboswitch, PK5 Mutant | 1 mM ZMP, cotranscriptional | Supp. Fig. 4 |

**Supplementary Table 3.**
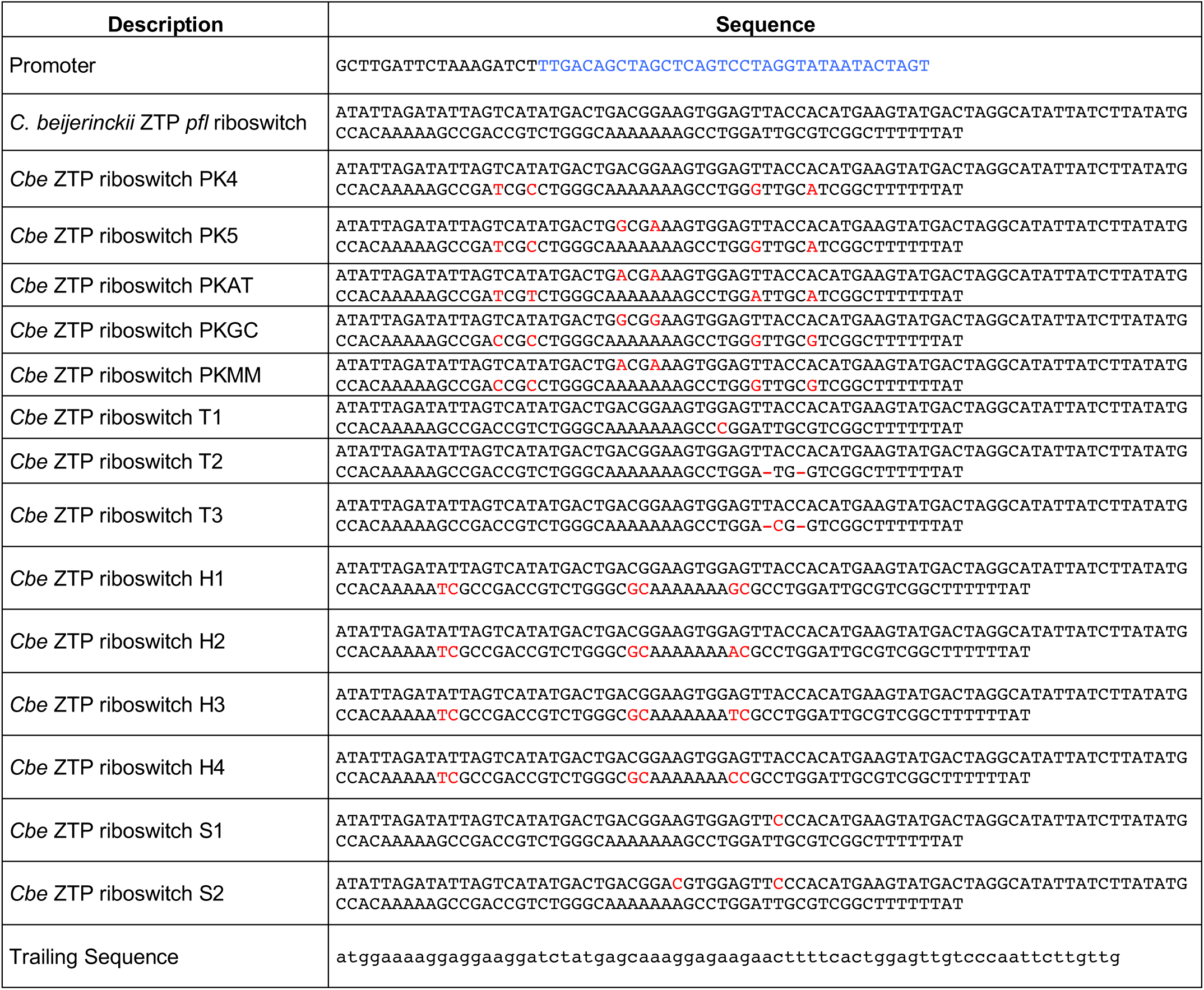
Sequences used for in vitro transcription templates. Summary of the sequences used for generating *in vitro* transcription templates. The ZTP riboswitch contains a two nucleotide insertion at the 5’ end of the template relative to the wild type *Clostridium beijerinckii* sequence. Templates were extended with a ribosome binding site (RBS) and a portion of the superfolder GFP (SFGFP) coding sequence, listed as ‘trailing sequence’ below. Promoter sequences are blue. Mutation positions are highlighted with red. Deletion positions are shown as “-”.

**Supplementary Table S4.**
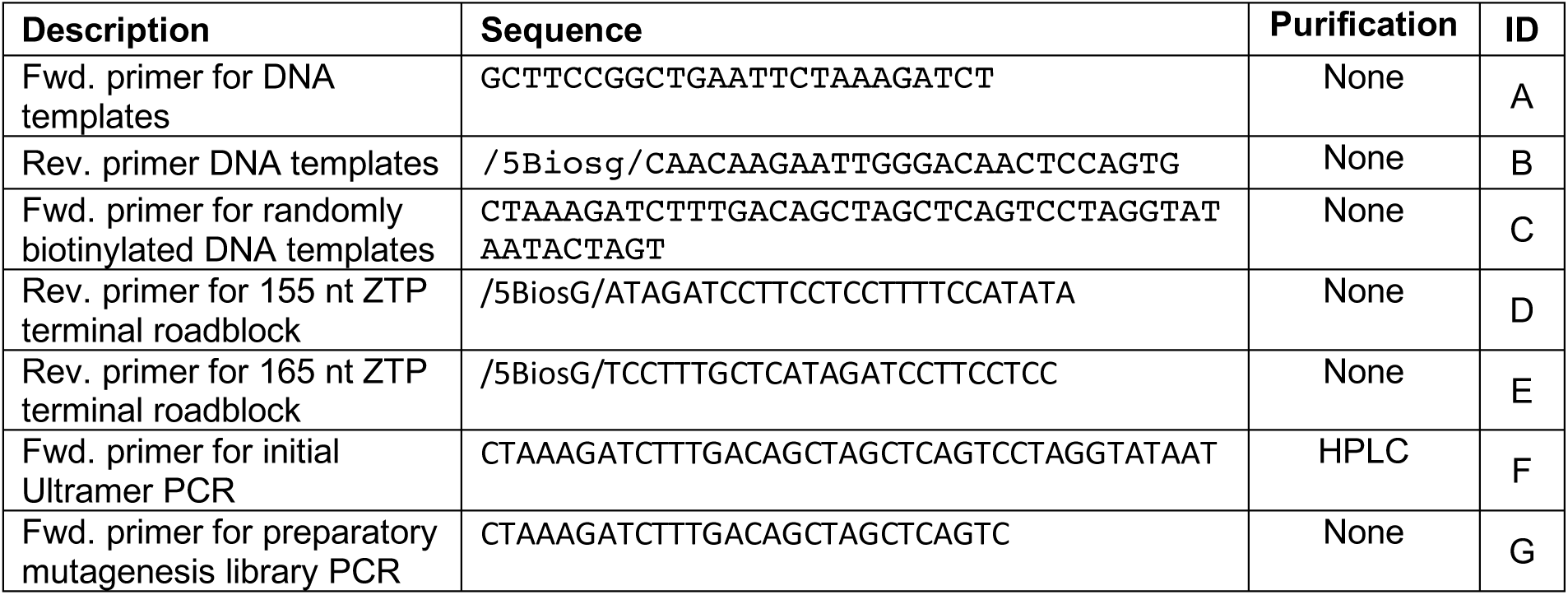
Oligonucleotides used for DNA template amplification. Below is a table of oligonucleotides used for the preparation of *in vitro* transcription DNA templates. Abbreviations within primer sequences are as follows: ‘/5Biosg/’ is a 5’ biotin moiety. These abbreviations were used for compatibility with the Integrated DNA Technologies ordering notation.

**Supplementary Table S5.**
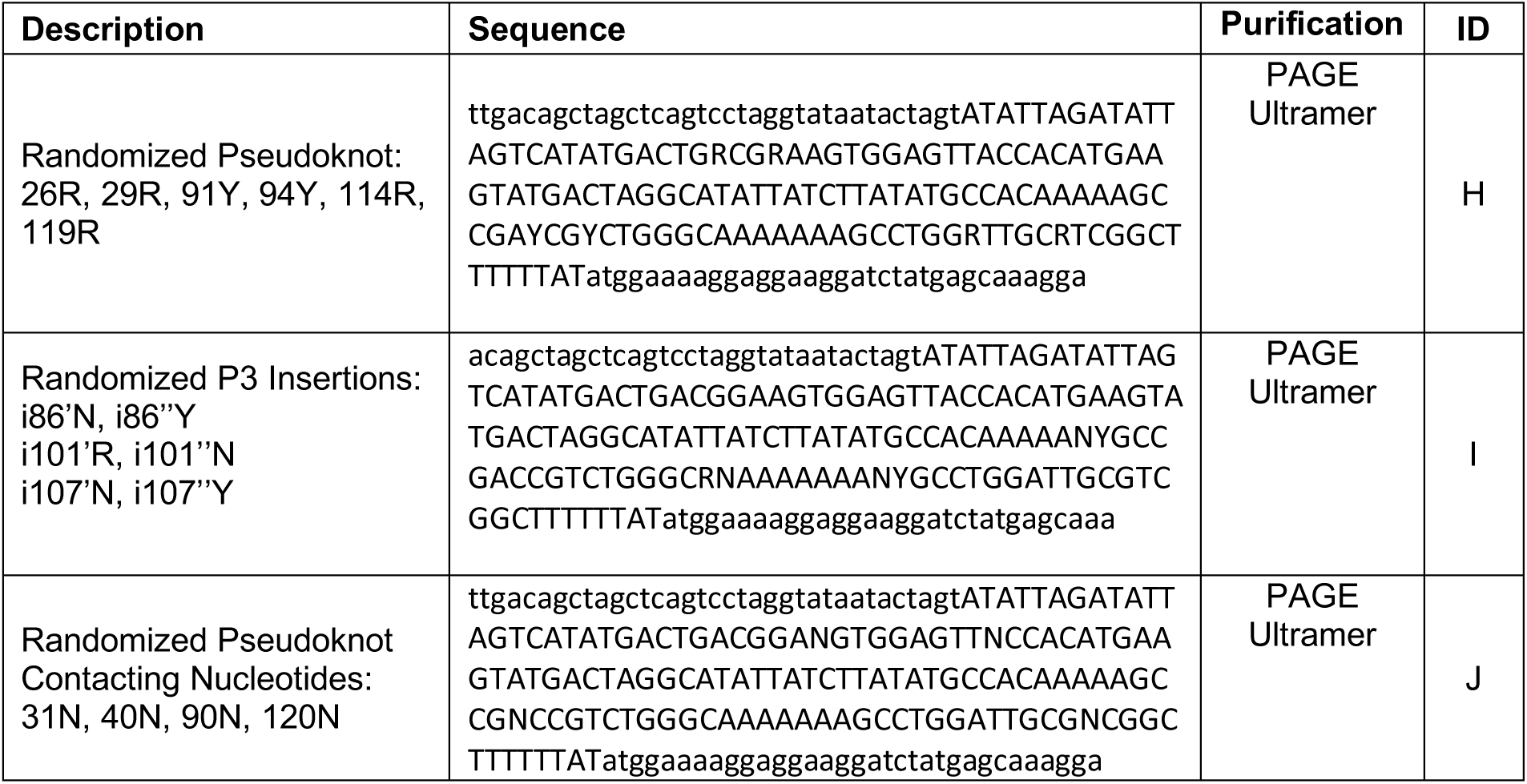
Ultramer Mutagenesis Oligonucleotides. Below is a table of Ultramer oligonucleotides used for the preparation of *in vitro* transcription DNA templates for combinatorial functional mutagenesis experiments. Uppercase sequence indicates the riboswitch. Lowercase sequence indicates flanking regions. IUPAC nucleotide notation is used. Insertion mutations are designated in the description by ‘i’ followed by the nucleotide number that that the insertion precedes; multiple insertions at a given position are designated by apostrophes. For example, NY insertions at nucleotide 86 are designated i86’N for the first insertion and i86’’Y for the second insertion.

**Supplementary Table 6.**
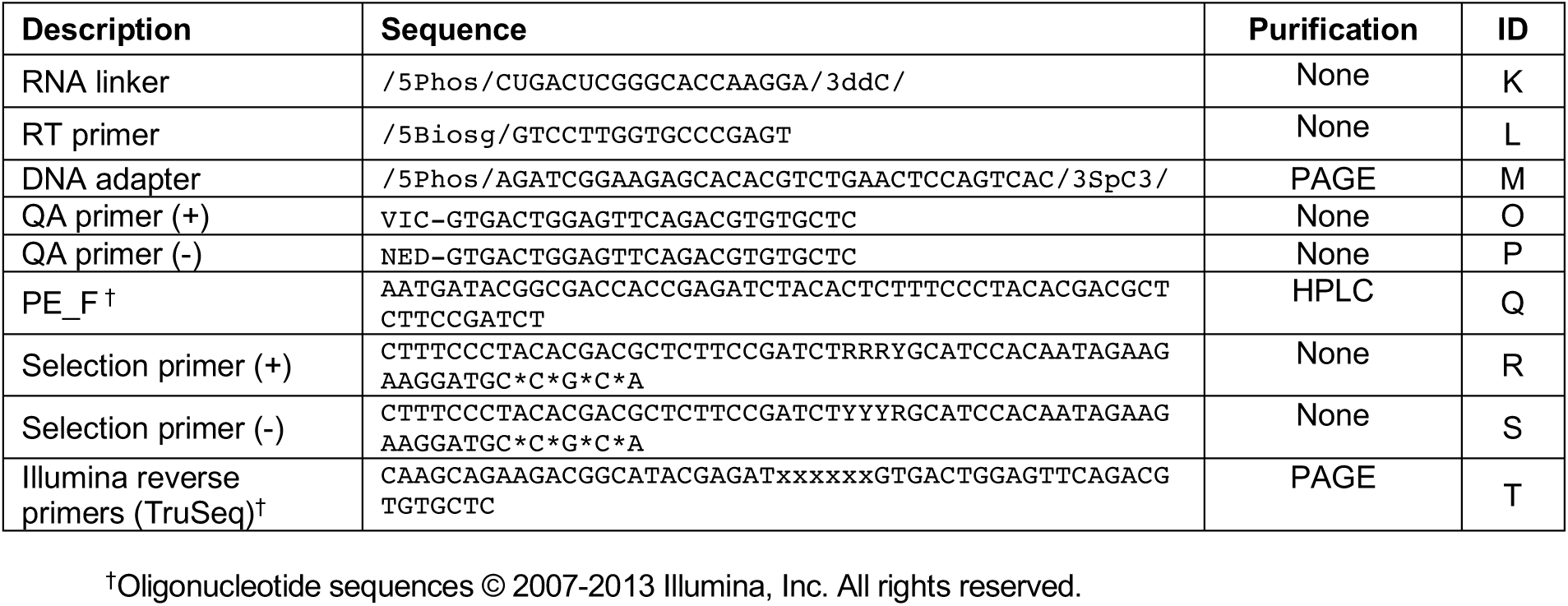
Oligonucleotides used for sequencing library preparation. Below is a table of oligonucleotides used during the cotranscriptional SHAPE-Seq and combinatorial mutagenesis experiments. The ‘xxxxxx’ sequence in the Illumina primer represents any TruSeq index. For the reverse template amplification primers, the sequence ‘NNNNNN…’ is the reverse complement from the 3’ end of the intermediate length being amplified. Abbreviations within primer sequences are as follows: ‘/5Biosg/’ is a 5’ biotin moiety, ‘/5Phos/’ is a 5’ monophosphate group, ‘/3SpC3/’ is a 3’ 3-carbon spacer group, /3ddC/ is a 3’ di-deoxy CTP, VIC and NED are fluorophores (ABI), and asterisks indicate a phosphorothioate backbone modification. These abbreviations were used for compatibility with the Integrated DNA Technologies ordering notation.

